# Male cuticular pheromones stimulate removal of the mating plug and promote re-mating through pC1 neurons in *Drosophila* females

**DOI:** 10.1101/2023.11.28.568981

**Authors:** Minsik Yun, Do-Hyoung Kim, Tal Soo Ha, Kang-Min Lee, Eungyu Park, Markus Knaden, Bill S. Hansson, Young-Joon Kim

## Abstract

In birds and insects, the female uptakes sperm for a specific duration post-copulation known as the ejaculate holding period (EHP) before expelling unused sperm and the mating plug through sperm ejection. In this study, we found that *Drosophila melanogaster* females shortens the EHP when incubated with males or mated females shortly after the first mating. This phenomenon, which we termed <u>m</u>ale-induced <u>E</u>HP <u>s</u>hortening (MIES), requires Or47b+ olfactory and ppk23+ gustatory neurons, activated by 2-methyltetracosane and 7-tricosene, respectively. These odorants raise cAMP levels in pC1 neurons, responsible for processing male courtship cues and regulating female mating receptivity. Elevated cAMP levels in pC1 neurons reduce EHP and reinstate their responsiveness to male courtship cues, promoting re-mating with faster sperm ejection. This study established MIES as a genetically tractable model of sexual plasticity with a conserved neural mechanism.

**Significance Statement:** Sexual plasticity, the adaptation of reproductive behavior to social changes, was explored in the fruit fly, a genetically tractable model insect. Our findings revealed that inseminated females, encountering another courting male post-mating, shorten the ejaculate holding period (EHP). Specific olfactory and gustatory pathways regulating this phenomenon were identified, converging on the pC1 neurons in the brain-a conserved neural circuit that regulates female mating activity. Odors associated with EHP shortening increased the second messenger cAMP. The transient elevation of cAMP heightened the excitability of pC1 neurons, facilitating the prompt removal of the male ejaculate and subsequent re-mating . This study established a behavioral model of sexual plasticity and provided a framework for understanding the neural circuit processes involved.

## Introduction

Sexual plasticity, the ability to modify sexual state or reproductive behavior in response to changing social conditions, is observed in both vertebrates and invertebrates (1–5). In rodents, exposure to unfamiliar males often leads to the sudden termination of pregnancy, known as the Bruce effect. It is induced by male urinary peptides, such as MHC I peptides, activating the vomeronasal organ (6–8). This effect enhances reproductive fitness of both sexes, by eliminating the offspring of competing males and enabling females to select better mates even after conception. Many species also adapt their reproductive behavior in response to the social sexual context change (SSCC), involving encounters with new sexual partners or competitors. Understanding the neural circuit mechanisms behind female responses to SSCC emerges as a central focus of neuroscience (9–12).

*Drosophila melanogaster,* the fruit fly, displays various social behaviors like aggregation, aggression, and sexual behavior (13–15). Similar to rodents, they primarily use the olfactory system to communicate socially through pheromones (16, 17). Some of these pheromones act as aphrodisiacs, while others regulate aggression or foster aggregation. For instance, cis-vaccenyl acetate (cVA) attracts females but repels males and promotes aggregation (14, 18, 19). 7-Tricosene (7-T), a cuticular hydrocarbon (CHC) present in males, is an aphrodisiac to females and affects social interactions between males (20, 21). On the other hand, 7,11-heptacosadiene (7,11-HD), a related female-specific pheromone, functions as an aphrodisiac to males, triggering courtship behavior and involving species recognition (22, 23).

The fruit fly’s chemo-sensory organs, located in various parts of the body, detect these pheromones (24, 25). Olfactory receptor neurons (ORNs) in the sensilla of the antennae and maxillary palp are responsible for the detection of long-range volatile pheromones like cVA, while short-range pheromones like 7-T are sensed by neurons on the fore-legs and labellum (16, 17, 24).

The olfactory receptor Or47b, expressed ORNs located in at4 trichoid sensilla on the third antennal segment, is involved in several socio-sexual interactions, including male mating success, mate preference, and female aggression toward mating pairs (12, 26–29). In males, Or47b senses fatty acid methyl esters and fatty acids that affect mating competition and copulation (30, 31). While the role of Or47b in female aggression is well established (12), its involvement in female sexual behavior is uncertain. In both sexes, Or47b ORNs project to VA1v glomeruli, where VA1v projection neurons receive their signal and project to the mushroom body calyx and lateral horn. Male Or47b neurons connect to neurons such as aSP5, aSP8, and aSP9, which express a male-specific transcription factor Fru^M^ (32).

CHC pheromones, which function as short-range pheromones, are detected primarily by neurons on the fore-legs and the labellum that express gustatory receptors (GR), ionotropic receptors (IR), or the ppk/DEG-ENaC family of sodium channels (16, 17, 24). CHCs like 7-T and 7,11-HD are sensed by *ppk23*-expressing M and F cells in the tarsi (33). 7-T and cVA are sensed by M cells expressing *ppk23*, whereas 7,11-HD and 7,11-nonacosadiene (7,11-ND) are sensed by F cells expressing *ppk23*, *ppk25*, and *ppk29*. In males 7-T or 7,11-HD affects the neuronal activity of the Fru^M^-expressing P1 neurons (34–36). However, how these CHCs signal in the female brain remains unknown.

Sperm ejection is a process by which females can remove the male ejaculate or the mating plug after copulation. This phenomenon has been observed in several animal species including feral fowl (37), black-legged kittiwake (38), and dunnock (39). In the fruit fly, it typically occurs approximately 90 minutes after mating (40). This specific interval, referred to as ejaculate holding period (EHP), is thought to affect sperm usage and fecundity (40, 41). The neurosecretory neurons in the brain pars intercerebralis (PI) that produce diuretic hormone 44 (Dh44), an insect orthologue of the corticotropin-releasing factor, regulate EHP (40). There is evidence that *Drosophila* females sense the social-sexual context through sperm ejected by other females. For instance, females were likely to lay more eggs when placed on a food patch containing male ejaculate deposited by other females (42). However, it remains unknown whether the SSCC influences sperm ejection and EHP.

Female pC1 neurons, which express a specific transcription factor Dsx^F^, integrate olfactory and auditory cues associated with male courtship (43, 44). The pC1 neurons, their male counterparts (i.e., P1 neurons), and the ventrolateral subdivision of ventromedial hypothalamus (VMHvl) neurons in mice share conserved circuit configurations and demonstrate functional similarity in coordinating social and sexual behaviors (45, 46). There are 14 Dsx-positive pC1 neurons in each hemisphere of the brain, responsive to the male sex-pheromone cVA and courtship songs (44, 47). Connectome analyses identified 10 pC1 neurons that fall into five subtypes, with pC1a, b, and c subtypes associated with mating behavior and pC1d and e subtypes associated with aggression (47–52). Although direct evidence connecting pC1 neurons to sperm ejection is limited, they are promising candidates for regulating sperm ejection or EHP, because sperm ejection allows females to eliminate the mating plug and male ejaculate, thereby restoring sexual attractiveness (53).

In this study, we demonstrated that two male pheromones, 2-methyltetracosane (2MC) and 7-T, significantly reduced the EHP through *Or47b* neurons and *ppk23* neurons, respectively. These pheromone pathways converge on pC1 neurons, where they increase cAMP levels. The elevated cAMP in pC1 neurons resulted in a reduction of the EHP to a degree that was comparable to the effects of the male pheromones. It also enhanced the excitability of pC1 neurons, making them more responsive to both olfactory and auditory male courtship cues and promoting further mating following the earlier removal of the mating plug. These findings establish a novel behavioral paradigm that sheds light on the intricate molecular and neuronal pathways underlying female sexual plasticity.

## Results

### Male-induced EHP shortening (MIES) is dependent on olfaction

To investigate the impact of changes in the social sexual context on the EHP, we compared the EHP of post-mating females isolated from any male presence to those exposed to naive wild-type *Canton-S* (*CS*) males immediately after copulation (Fig. 1A). Notably, the EHP of females incubated with naive males was approximately 30 minutes shorter than that of females left in isolation after mating (Fig. 1A, 1B). We refer to this phenomenon as male-induced EHP shortening (MIES). In contrast, little difference in EHP was observed between females incubated with virgin females and those isolated after mating (Fig. 1C).

**Fig. 1.**
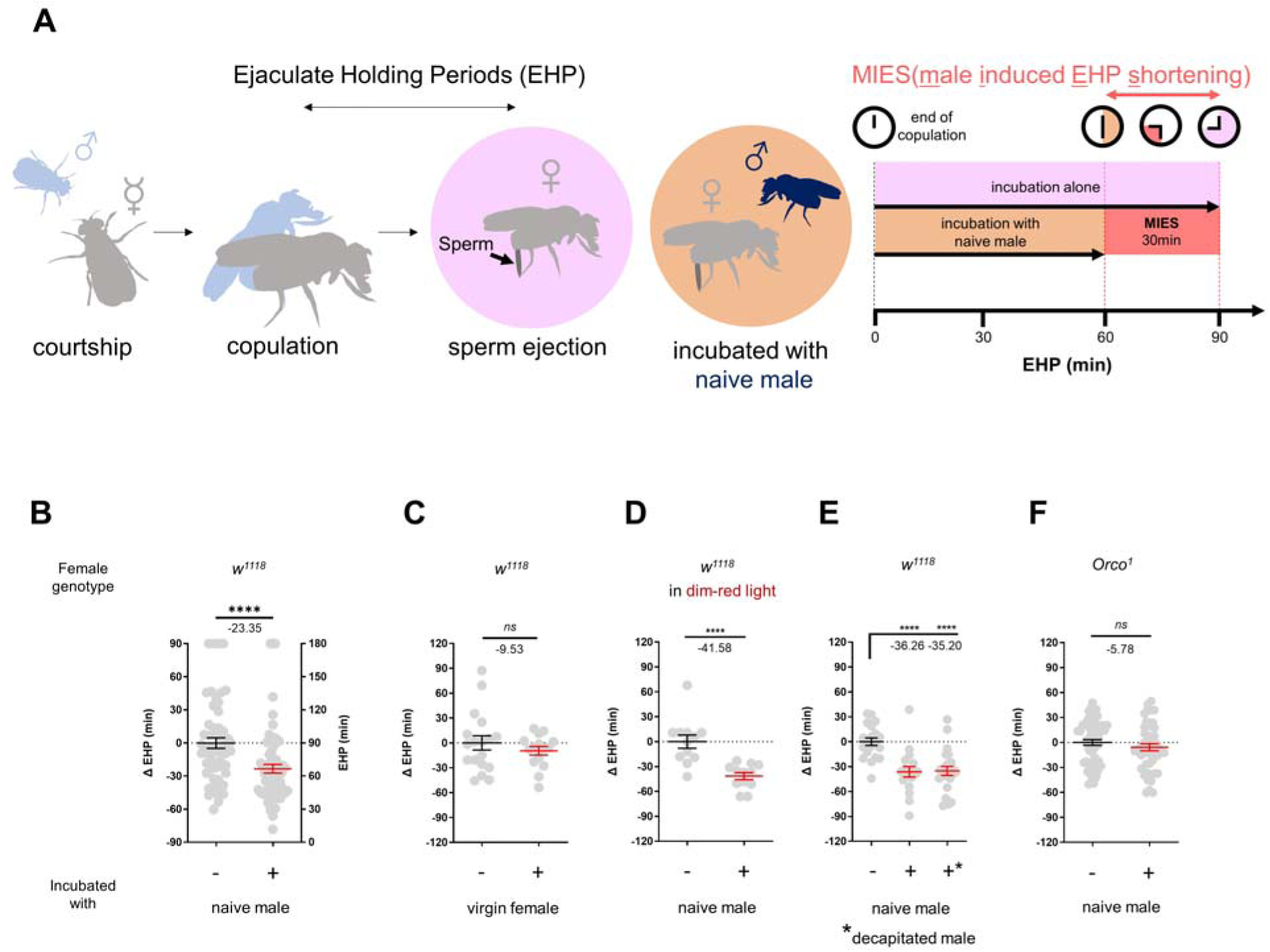
The presence of males reduces the ejaculate holding period (EHP) in females through olfactory or gustatory sensation. **A**, Schematic of the experimental procedure employed to measure male-induced EHP shortening (MIES). Immediately after the end of copulation, the female is incubated with a wild-type *Canton-S (CS)* male that has not been previously exposed to the female. Typically, *w^1118^*females that are kept alone after mating exhibit an EHP of approximately 90 minutes, whereas females that are incubated with a naïve *CS* male exhibit an EHP of approximately 60 minutes. In this study, we refer to this phenomenon as MIES. **B-F**, Normalized ejaculate holding period (EHP) or ΔEHP of the females of the indicated genotypes, incubated under the indicated conditions after mating. The ΔEHP is calculated by subtracting the mean of the reference EHP of females kept alone after mating (the leftmost column) from the EHP of individual females in comparison. Mann-Whitney Test (n.s. *p* > 0.05; \*\*\*\**p* < 0.0001). Gray circles indicate the EHP or ΔEHP of individual females, and the mean ± SEM of data is presented. Numbers below the horizontal bar represent the mean of the EHP differences between the indicated treatments.

Male fruit flies employ various sensory signals to attract females during courtship (15). To assess the role of the visual signal in MIES, we examined MIES under dim red light conditions and observed that limited illumination had a marginal impact on MIES (Fig. 1D). Next, we examined MIES in post-mating females incubated with decapitated *CS* males. These males could serve as a source of olfactory or gustatory signals, but not for auditory or visual signals. Again, no reduction in MIES was observed (Fig. 1E). This strongly suggests that olfactory or gustatory cues are the key signals responsible for MIES. This is further supported by the observation that females deficient in the odorant receptor co-receptor (*Orco*^1^) did not exhibit MIES (Fig. 1F). Thus, it is highly likely that male odorant(s), especially those detected by olfactory receptors (Or), induce MIES.

### MIES is dependent on the *Or47b* receptor and *Or47b*-expressing ORNs

In the fruit fly antenna, the trichoid sensilla and their associated olfactory receptor neurons are known to detect sex pheromones (54). To investigate the contribution of ORNs located in the trichoid sensilla to MIES, we silenced 11 different ORN groups found in the trichoid and intermediate sensilla (55, 56) by expressing either the active or inactive form of Tetanus toxin light chain (TNT) (57). Our results showed that silencing ORNs expressing *Or13a*, *Or19a*, *Or23a*, *Or47b*, *Or65c*, *Or67d* or *Or88a* significantly affected MIES (Fig. S1A).

We then focused on the analysis of *Or47b*-positive ORNs (Fig. 2A), which, in contrast to the others, exhibited almost complete abolition of MIES when silenced. Activation of these neurons with the thermogenetic activator dTRPA1 (58) resulted in a significant EHP shortening, even in the absence of male exposure (Fig. 2B). Subsequently, we examined whether restoring Orco expression in *Or47b* ORNs in Orco-deficient females would restore MIES. Our results confirmed that this is indeed the case (Fig. 2C). To establish the necessity of the *Or47b* receptor gene for MIES, we examined Or47b-deficient females (*Or47b^2^/Or47b^3^*) and observed a complete absence of MIES, whereas heterozygous controls exhibited normal MIES (Fig. 2D). Furthermore, the reintroduction of *Or47b* expression in *Or47b* ORNs of *Or47b*-deficient females almost completely restored MIES (Fig. 2E). Based on these observations, we concluded that MIES depends on the *Or47b* receptor gene and Or47b-expressing ORNs.

**Fig. 2.**
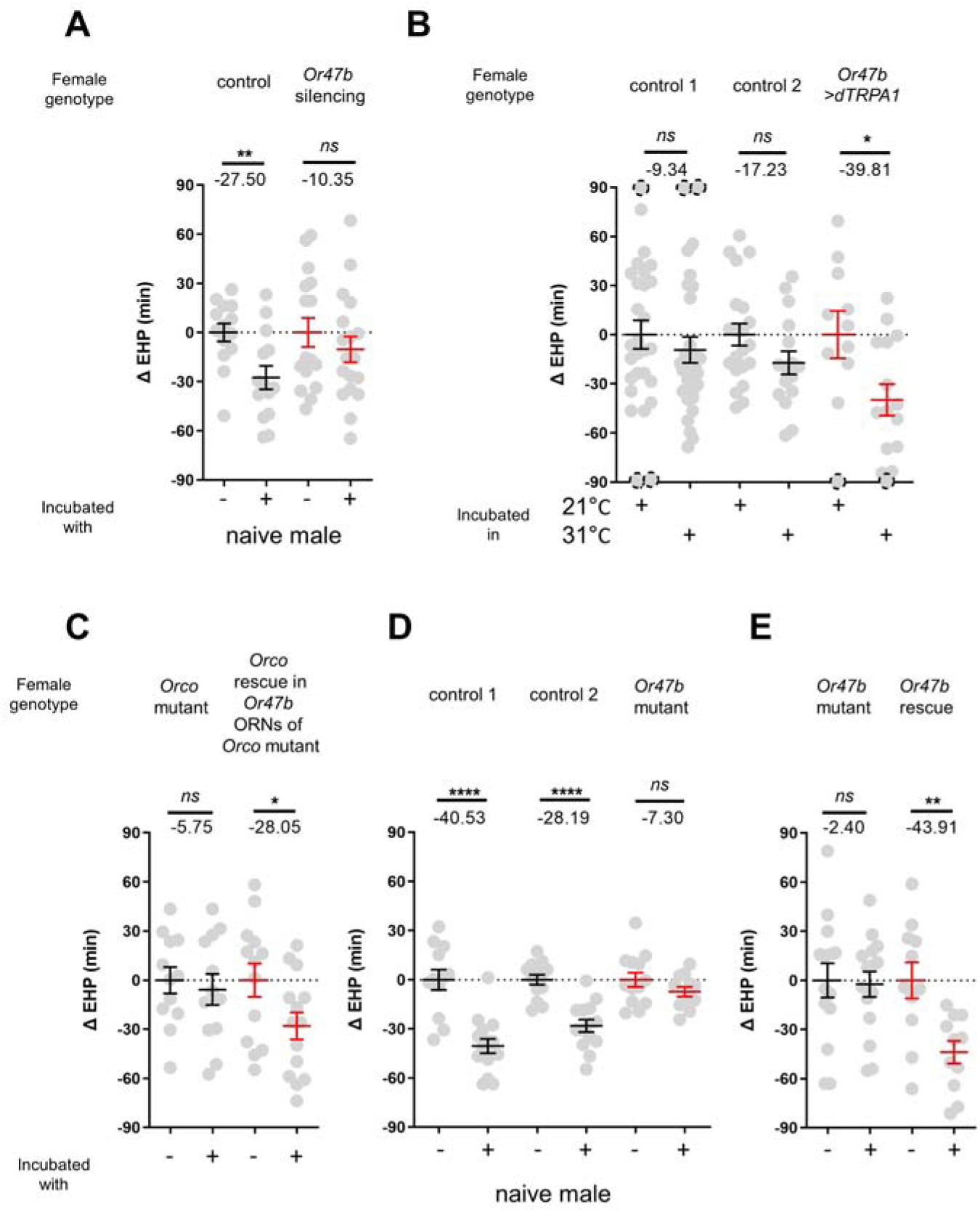
The function of Or47b and Or47b-positive ORNs is essential for MIES. **A, C-E**, ΔEHP of females of the indicated genotypes, incubated with or without naive males after mating. The female genotypes are as follows from left to right: (A) control (*Or47b>TNT^inactive^*), *Or47b* ORN silencing (*Or47b>TNT^active^*); (C) *Orco* mutant (*Orco^1^*/*Orco^1^*), *Orco* rescue in *Or47b* ORNs of *Orco* mutant (*Orco^1^*/*Orco^1^*; *Or47b>Orco*); (D) control 1 (*Or47b^2^/+*), control 2 (*Or47b^3^/+*), *Or47b* mutant (*Or47b^2^/Or47b^3^*); (E) *Or47b* mutant *(Or47b^2^/Or47b^2^*), *Or47b* rescue (*Or47b>Or47b; Or47b^2^/Or47b^2^*). **B**, Thermogenetic activation of *Or47b*-positive ORNs shortens EHP in females kept alone after mating. The female genotypes are as follows from left to right: control 1 (*Or47b-Gal4/+),* control 2 (*UAS-dTRPA1/+), Or47b>dTRPA1 (Or47b-Gal4/UAS-dTRPA1*). Mann-Whitney Test (n.s. *p* > 0.05; \**p* <0.05; \*\**p* <0.01; \*\*\*\**p* < 0.0001). The ΔEHP is calculated by subtracting the mean of the reference EHP of females kept alone after mating (‘-’ in A, C-E) or incubated at 21°C control conditions (B) from the EHP of individual females in comparison. Gray circles indicate the ΔEHP of individual females, and the mean ± SEM of data is presented. The gray circles with dashed borders indicate ΔEHP values that exceed the axis limits (>90 or <-90 minutes). Numbers below the horizontal bar represent the mean of the EHP differences between the indicated treatments.

### 2-Methyltetracosane (2MC) induces MIES via *Or47b* and *Or47b* ORNs

Previous studies have shown that methyl laurate (ML) and palmitoleic acid (PA) can activate *Or47b* ORNs only in the presence of a functional *Or47b* gene (30, 31). However, in our investigation, none of these odorants induced significant EHP shortening, even when applied at concentrations as high as 1440 ng (Fig. S2). This prompted us to search for a new pheromone capable of activating Or47b ORNs and thereby shortening the EHP.

Oenocytes produce a significant portion of the cuticular hydrocarbons or pheromones. We asked whether the male pheromone responsible for MIES is produced by oenocytes (Fig. S3A). Indeed, incubation with females engineered to produce male oenocytes significantly shorten EHP, strongly suggesting that male oenocytes serve as a source for the MIES pheromone. Unexpectedly, however, incubation with males possessing feminized oenocytes also resulted in significant EHP shortening (Fig. S3A). This raises the possibility that oenocytes may not be the sole source of the MIES pheromone, implying the involvement of more than one pheromone, for instance one from oenocytes and another from an alternate source, in MIES.

The genus *Drosophila* exhibits distinct CHC profiles, with certain CHC components shared among closely related species (59). We found that incubation with males of other closely related species, such as *D. simulans*, *D. sechellia*, and *D. erecta,* also induced EHP shortening, whereas incubation with *D. yakuba* males did not (Fig. S3B).

The EHP was therefore measured in females incubated in a small mating chamber containing a piece of filter paper perfumed with male CHCs, including 2-methylhexacosane, 2-methyldocosane, 5-methyltricosane, 7-methyltricosane, 10Z-heneicosene, 9Z-heneicosene, and 2MC at various concentrations (not shown). Among these, 2MC at 750 ng was the only one that significantly reduced EHP (Fig. 3A; Fig. S4). 2MC was mainly found in males, but not in virgin females (30). Notably, it is present in *D. melanogaster*, *D. simulans*, *D. sechellia*, and *D. erecta*, but not in *D. yakuba* (30, 60).

**Fig. 3.**
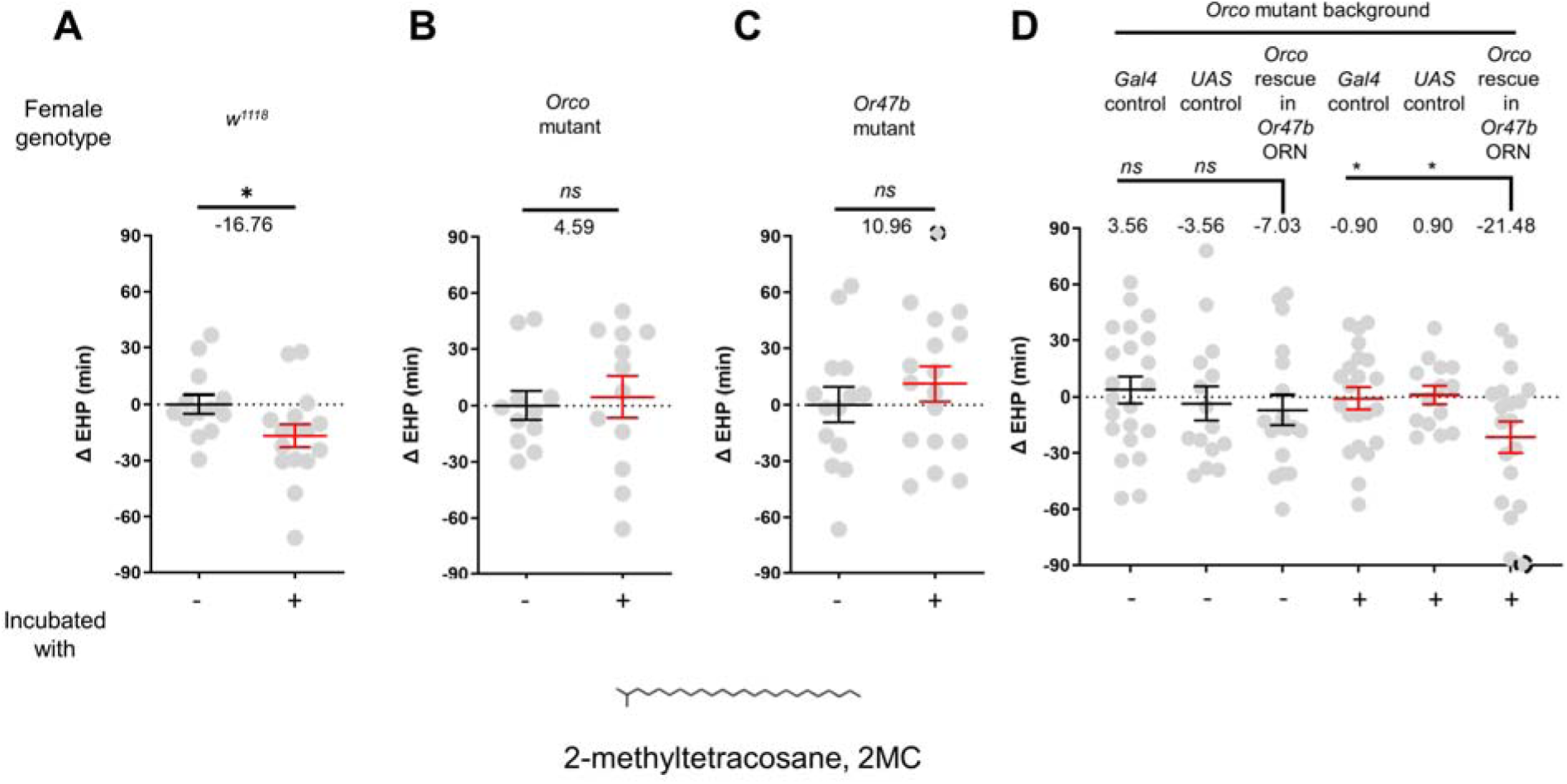
2-Methyltetracosane (2MC) can induce EHP shortening through Or47b. **A-D**, Δ EHP of females of the indicated genotypes, incubated in solvent vehicle or 2MC. Mated females were incubated with a piece of filter paper perfumed with either vehicle (-) or 750 ng 2MC (+). The female genotypes are as follows: (A) *w^1118^*, (B) *Orco* mutant *(Orco^1^*/*Orco^1^*), (C) *Or47b* mutant *(Or47b^2^/Or47b^2^*), (D) *Gal4* control (*Or47b-Gal4/*+; *Orco^1^*/*Orco^1^*), *UAS* control (*UAS-Orco/+*; *Orco^1^*/*Orco^1^*), *Orco* rescue in *Or47b* ORN (*Orco^1^*/*Orco^1^*; *Or47b-Gal4/UAS-Orco*). A-C , Mann-Whitney Test (n.s. *p* > 0.05; \**p* <0.05). D, One-way analysis of variance (ANOVA) test with Fisher’s LSD multiple comparison (n.s. *p* > 0.05; \**p* < 0.05). Gray circles indicate the ΔEHP of individual females and the mean ± SEM of data is presented. The ΔEHP is calculated by subtracting the mean of the reference EHP of females incubated with vehicle-perfumed paper (the leftmost column in A-C) or the mean of the *Gal4* control and *UAS* control female incubated with vehicle-perfumed paper (the two leftmost columns in D) from the EHP of individual females in comparison. Gray circles with dashed borders indicate ΔEHP values that exceed the axis limits (>90 or <-90 minutes). Numbers below the horizontal bar represent the mean of the EHP differences between the indicated treatments.

Moreover, the 2MC-induced EHP shortening was not observed in Orco- or Or47b-deficient females (Fig. 3B, 3C), but was restored when Orco expression was reinstated in *Or47b* ORNs in Orco-deficient mutants (Fig. 3D). Our behavioral observations strongly suggest that 2MC acts as an odorant ligand for Or47b and shortens the EHP through this receptor.

### 7-Tricosene (7-T) shortens EHP through *ppk23* neurons

In contrast to incubation with virgin females, incubation with mated females resulted in a significant shortening of EHP (Fig. 4A). Mated females carry male pheromones, including 7-T and cVA, which are transferred during copulation (53). This raised the possibility that these male pheromones might also induce EHP shortening. Indeed, our experiments revealed that incubation with a piece of filter paper perfumed with 150 ng of 7-T significantly shortened the EHP. Conversely, incubation with cVA and 7-pentacosene, a related CHC, did not produce the same effect (Fig. 4B, 4C; Fig. S5A-5B). The concentrations of 7-T capable of inducing EHP shortening appear to be physiologically relevant. 7-T has been found at levels of 432 ng in males (61), 25 ng in virgin females, and 150 ng in mated females (53). Although the receptors for 7-T remain unknown, *ppk23*-expressing tarsal neurons have been shown to sense these compounds and regulate sexual behavior in males and females (23, 62, 63). Subsequently, we silenced *ppk23* neurons, and as a result, MIES was almost completely abolished, underscoring the pivotal role of 7-T in MIES (Fig. 4D). However, DEG/ENac channel genes expressed in *ppk23* neurons, including *ppk23* and *ppk29*, were found to be dispensable for MIES (Fig. S5C-5E). This aligns with previous observations that neither *ppk23* deficiency nor *ppk28* deficiency recapitulates the sexual behavioral defects caused by silencing *ppk23* neurons (64).

**Fig. 4.**
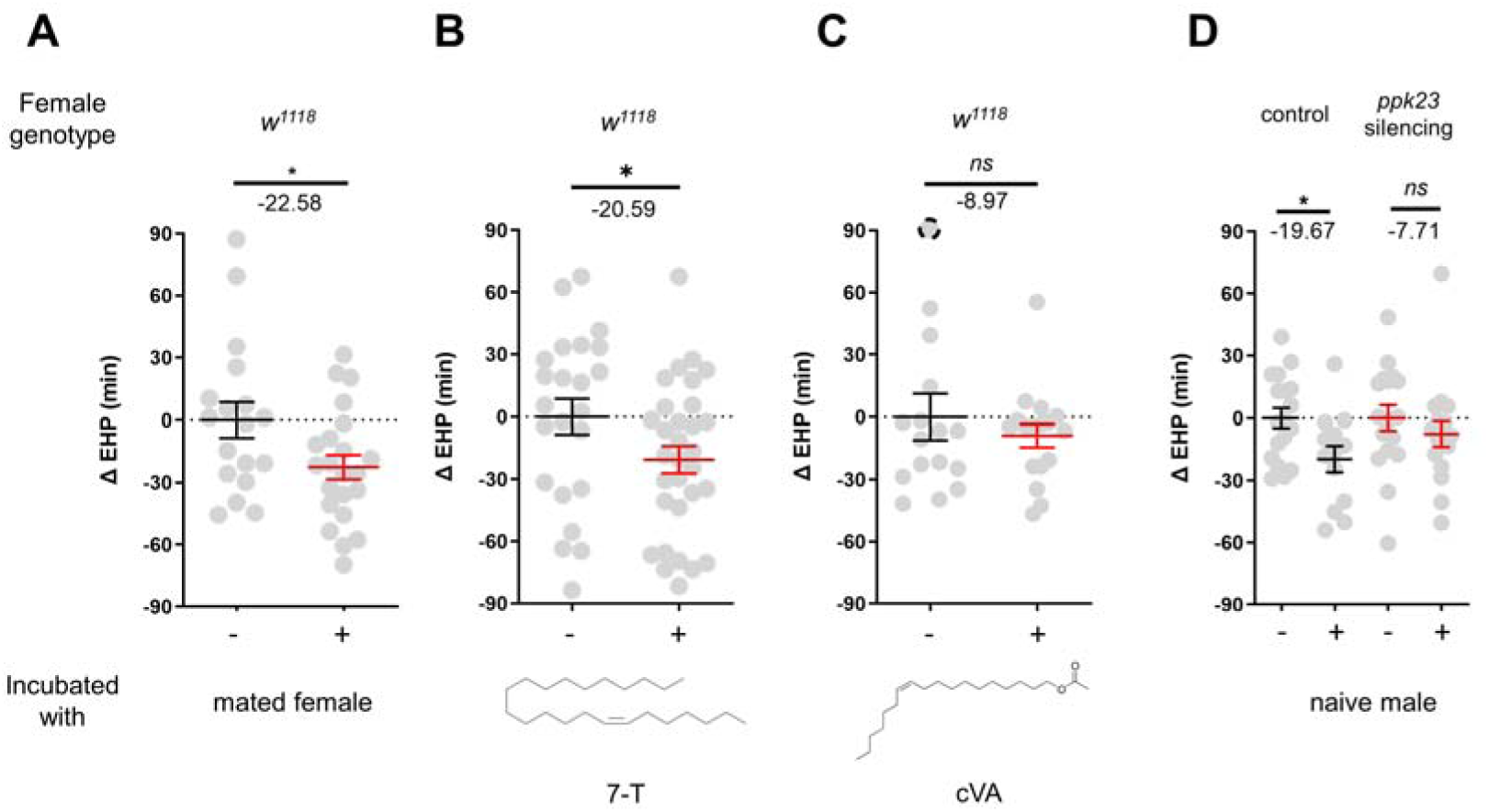
7-Tricosene present in mated females and males reduces EHP via *ppk23* neurons. A-D , ΔEHP of females of the indicated genotypes, incubated with mated females (A), a piece of filter paper perfumed with 150 ng 7-T (B), 200 ng cVA (C), or naive males (D) after mating. The female genotypes are as follows: (A-C) *w^1118^*, (D) control (*ppk23-Gal4/UAS-TNT^inactive^*), *ppk23* silencing (*ppk23-Gal4/UAS-TNT^active^*). A, Unpaired *t*-Test, B-D, Mann-Whitney Test (n.s. *p* > 0.05; \**p* < 0.05). The ΔEHP is calculated by subtracting the mean of the reference EHP of females kept alone (‘-’ in A, D) or incubated with vehicle-perfumed paper (the leftmost column in B, C) from the EHP of individual females in comparison. Gray circles indicate the ΔEHP of individual females, and the mean ± SEM of data is presented. The gray circles with dashed borders indicate ΔEHP values that exceed the axis limits (>90 or <-90 minutes). Numbers below the horizontal bar represent the mean of the EHP differences between the indicated treatments.

### The pC1 b and c neurons regulate EHP and MIES

The neuropeptide Dh44 determines the timing of sperm ejection or EHP (40). The same study found that Dh44 receptor neurons involved in EHP regulation also express *doublesex* (*dsx*), which encodes sexually dimorphic transcription factors. A recent study has revealed that pC1 neurons, a specific subgroup of *dsx*-expressing central neurons in the female brain, do indeed express Dh44 receptors (47). With these findings, we set out to investigate the role of pC1 neurons in the regulation of EHP and MIES. The pC1 neurons comprise five distinct subtypes. Of these, the pC1a, b, and c subtypes have been implicated in mating receptivity (47, 48), while the remaining pC1d and e subtypes have been implicated in female aggression (48, 52). To investigate the role of these subtypes in EHP, we employed GtACR1, an anion channel activated by blue light in the presence of all *trans*-retinal (ATR), to silence specific pC1 subtypes immediately after mating. Our experiments revealed that silencing of the pC1 subset comprising the pC1a, b and c subtypes with GtACR1 led to an increase in EHP (Fig. 5A), whereas silencing of the pC1d and e subtypes had a limited effect on EHP (Fig. 5B). We further analyzed the roles of pC1b, c neurons along with pC1a neurons separately. We generated a subtype-specific split-Gal4 for pC1a and found that, as expected, silencing pC1a with this split-Gal4 almost completely suppressed mating receptivity (Fig. S6).

**Fig. 5.**
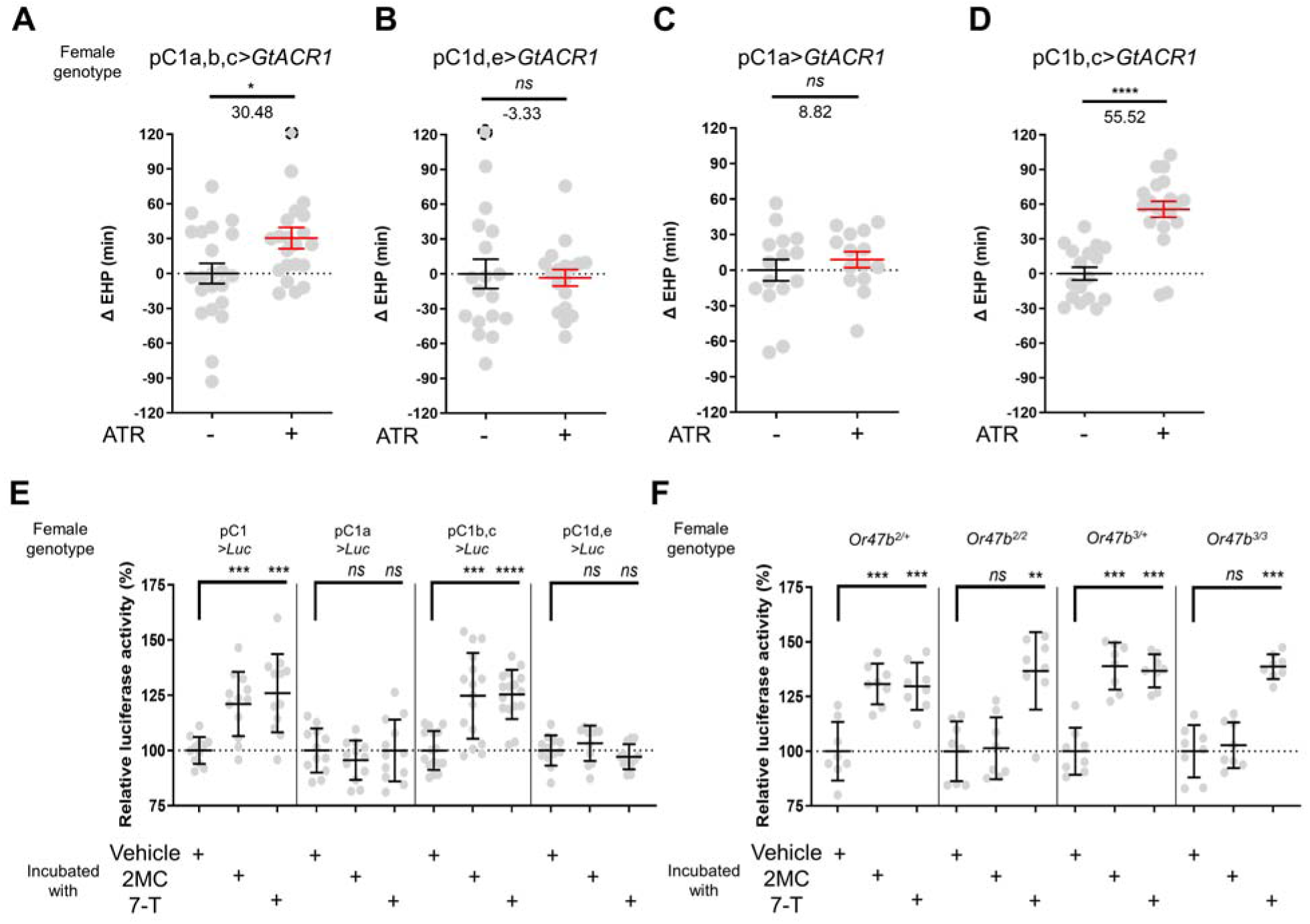
A subset of pC1 neurons, comprising pC1b and pC1c subtypes, regulates EHP and exhibits CRE-luciferase reporter activity in response to 2MC and 7-T. **A-D**, The optogenetic silencing of a pC1 neuron subset comprising pC1b and pC1c neurons (pC1b, c) increases EHP. Females of the indicated genotypes were cultured on food with or without all *trans*-retinal (ATR) after eclosion. The ΔEHP is calculated by subtracting the mean of the reference EHP of females cultured in control ATR-food (the leftmost column) from the EHP of individual females in comparison. The female genotypes are as follows: (A) pC1a,b,c*>GtACR1* (*pC1-S-Gal4/UAS-GtACR1*), (B) pC1d,e*>GtACR1* (*pC1-A-Gal4/UAS-GtACR1*), (C) pC1a*>GtACR1* (*pC1a-split-Gal4 /UAS-GtACR1*), and (D) pC1b,c*>GtACR1* (*Dh44-pC1-Gal4/UAS-GtACR1*). Gray circles indicate the ΔEHP of individual females, and the mean ± SEM of data is presented. The gray circles with dashed borders indicate ΔEHP values that exceed the axis limits (>120 minutes). Mann-Whitney Test (n.s. *p* > 0.05; \**p* <0.05; \*\*\*\**p* < 0.0001). Numbers below the horizontal bar represent the mean of the EHP differences between the indicated treatments. **E, F**, Relative CRE-luciferase reporter activity of pC1 neurons in females of the indicated genotypes, incubated with a piece of filter paper perfumed with solvent vehicle control or the indicated pheromones immediately after mating. The CRE-luciferase reporter activity of pC1 neurons of Or47b-deficient females (*Or47b^2/2^* or *Or47b^3/3^*) was observed to increase in response to 7-T but not to 2MC. To calculate the relative luciferase activity, the average luminescence unit values of the female incubated with the vehicle are set to 100%. Mann-Whitney Test (n.s. *p* > 0.05; \*\**p* < 0.01; \*\*\**p* < 0.001; \*\*\*\**p* < 0.0001). Gray circles indicate the relative luciferase activity (%) of individual females, and the mean ± SEM of data is presented.

However, silencing pC1a alone did not result in increased EHP, suggesting a marginal role of the pC1a subtype in EHP regulation (Fig. 5C). In contrast, concomitant silencing of both pC1b and pC1c neurons significantly increased EHP by 56 ± 6.9 minutes (Fig. 5D). At present, we lack the genetic tools to further distinguish the roles of pC1b and pC1c subtypes separately.

### 2MC and 7-T increase cAMP levels in pC1 b and c neurons

Our recent research has shown that pC1 neurons exhibit elevated cAMP levels during sexual maturation, and that this increase in cAMP is closely related to heightened excitability of pC1 neurons (47). The same study also showed that a mating signal (i.e., sex peptide in the male seminal fluid) reduces cAMP levels in pC1 neurons. Thus, we hypothesized that male odorants responsible for inducing MIES, such as 2MC or 7-T, would elevate cAMP levels in pC1b, c neurons in newly mated females. This, in turn, would lead to increased excitability of pC1 neurons and, as a consequence, a reduction in the EHP. To monitor cAMP levels in these neurons, we prepared females that express a CRE-luciferase reporter selectively in pC1b, c neurons. Indeed, when exposed to 2MC or 7-T, pC1b, c neurons exhibited a significant increase in CRE-luciferase activity, indicating that these neurons produce higher levels of cAMP in response to these odorants (Fig. 5E). Notably, CRE-luciferase activity appeared to peak at specific odorant concentrations that induced significant shortening of the EHP (Fig. S7).

In contrast, when we examined other pC1 subsets, such as pC1a, and pC1d and e, we detected no evidence of increased CRE-luciferase reporter activity upon exposure to 2MC or 7-T treatment (Fig. 5E). Notably, CRE-luciferase reporter activity in the pC1a neurons appears to be dependent on the mating status, as it reaches levels similar to those of pC1b, c neurons in virgin females (Fig. S8). This observation aligns well with connectome data suggesting that SAG neurons, which are responsible for relaying SP-dependent mating signals, synapse primarily with the pC1a subtype and to a much lesser extent with other pC1 subtypes (49).

To further test the role of Or47b in 2MC detection, we generated Or47b-deficient females with pC1 neurons expressing the CRE-luciferase reporter. Females with one copy of the wild-type Or47b allele, which served as the control group, showed robust CRE-luciferase reporter activity in response to either 2MC or 7-T. In contrast, *Or47b*-deficient females showed robust CRE-luciferase activity in response to to 7-T, but little activity in response to 2MC. This observation suggests that the odorant receptor Or47b plays an essential role in the selective detection of 2MC (Fig. 5F).

### Elevated cAMP in pC1 neurons shortens the EHP, while increasing re-mating

Having shown that MIES-inducing male odorants, 2MC or 7-T, increase cAMP levels in pC1b, c neurons from mated females, we next asked whether this induced elevation of cAMP levels in pC1b, c neurons would shorten EHP, leading to MIES. We employed the photoactivatable adenylate cyclase (PhotoAC), which increases cellular cAMP levels upon exposure to light. Indeed, the induced elevation of cAMP levels in pC1b, c neurons significantly shortened EHP, whereas the same treatment applied to pC1a or pC1d and pC1e had no such effect (Fig. 6A). This further underscores the pivotal role of pC1b, c neurons in EHP regulation.

**Fig. 6.**
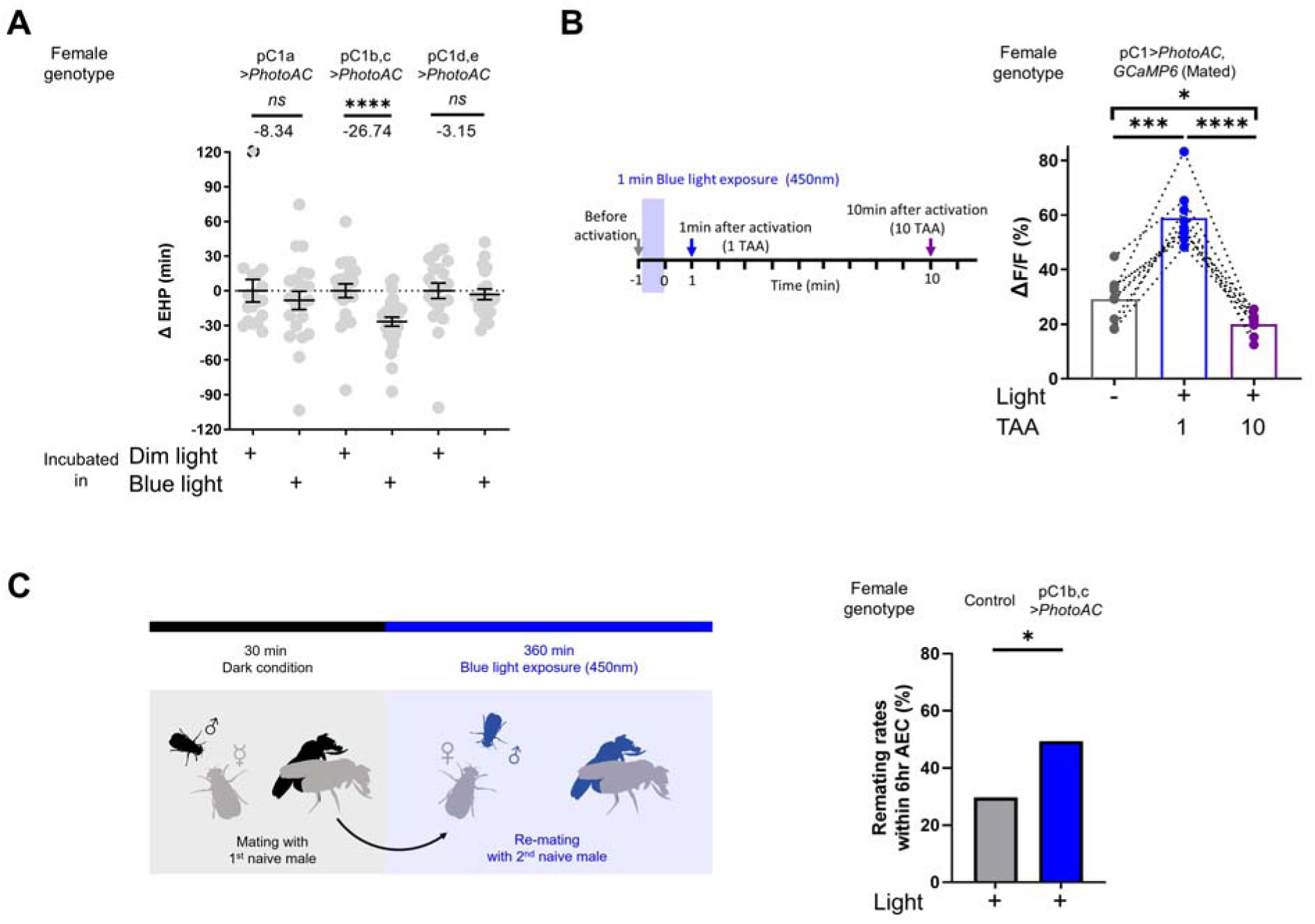
Elevated cAMP levels in pC1 neurons reduce EHP and increase the responsiveness of pC1 neurons to male courtship cues, thereby promoting subsequent remating. **A**, The optogenetic production of cAMP in the pC1b, c neurons shortens EHP, whereas the same treatment in pC1a or pC1d, e neurons does not. ΔEHP is calculated by subtracting the mean of the reference EHP of females incubated in the control illumination (Dim light), which does not activate a photoactivatable adenylate cyclase (PhotoAC), from the EHP of individual females. Mann-Whitney Test (n.s. *p* > 0.05, \*\*\*\**p* < 0.0001). **B**, The optogenetic production of cAMP transiently increases the excitability of pC1 neurons. Left, schematic of the experimental procedure. Right, peak ΔF/F in the LPC projections of pC1 neurons from freshly mated females in response to the pheromone cVA, before and after photoactivation of PhotoAC expressed in pC1 neurons. The calcium response was measured at specific time points after photoactivation: after 1 minute (blue dots and box) or 10 minutes (purple dots and box) after activation. Repeated measures one-way ANOVA test with the Geisser-Greenhouse correction followed by Tukey’s multiple comparisons test (\**p* < 0.05; \*\*\**p* < 0.001; \*\*\*\**p* < 0.0001). **C**, Left, schematic of the experimental procedure. Right, re-mating rate of females during optogenetic cAMP production in pC1b, c neurons, scored as the percentage of females that copulate with a naive *CS* male within 6 hours after the first mating. The female genotypes are as follows: Control (+/*UAS-PhotoAC*), pC1b,c*>UAS-PhotoAC* (*Dh44-pC1-Gal4/UAS-PhotoAC*). Chi-square test (\**p* <0.05).

Next, we asked whether the expression of Dh44R1 and Dh44R2, GPCRs that increase cellular cAMP in response to their ligand Dh44, in pC1b, c neurons is necessary for MIES. However, double knockdown of Dh44R1 and Dh44R2 in pC1 neurons seemed to have a limited impact on MIES (Fig. S9). This suggests that Dh44R signaling in pC1 neurons is not essential for the regulation of EHP or MIES, raising the possibility that other GPCRs may be involved in the up-regulation of cAMP levels in pC1 neurons in response to 2MC or 7-T.

Lastly, we investigated how increased cAMP levels affect the physiological activity of pC1 neurons. pC1 neurons from virgin females exhibit robust Ca^2+^ transients in response to male courtship cues, such as the male pheromone cVA and the courtship pulse song (44). In contrast, those from mated females display significantly diminished Ca^2+^ transients (47). When examined shortly after mating, a decrease in pC1 responsiveness to cVA was observed. However, immediately after PhotoAC activation in pC1 neurons, pC1 neurons from freshly mated females became more excitable and exhibited stronger Ca^2+^ transients in response to cVA (Fig. 6B). It is important to note that this PhotoAC-induced increase in pC1 excitability is transient and rapidly declines within 10 minutes (Fig. 6B). Nevertheless, these findings suggest that the increased cAMP levels in pC1 neurons would not only promote MIES but also facilitate re-mating in post-mating females, which typically engage in re-mating at a low frequency. To test this hypothesis, we examined the re-mating frequency of freshly mated females paired with naive males while inducing a cAMP increase in pC1 neurons. As expected, PhotoAC activation in pC1b, c neurons substantially increased the re-mating rate compared to the control group (Fig. 6C). Therefore, we concluded that male odorants that stimulate cAMP elevation in pC1 neurons expedite the removal of the mating plug, consequently leading to increased instances of re-mating (Fig. 7).

**Fig. 7.**
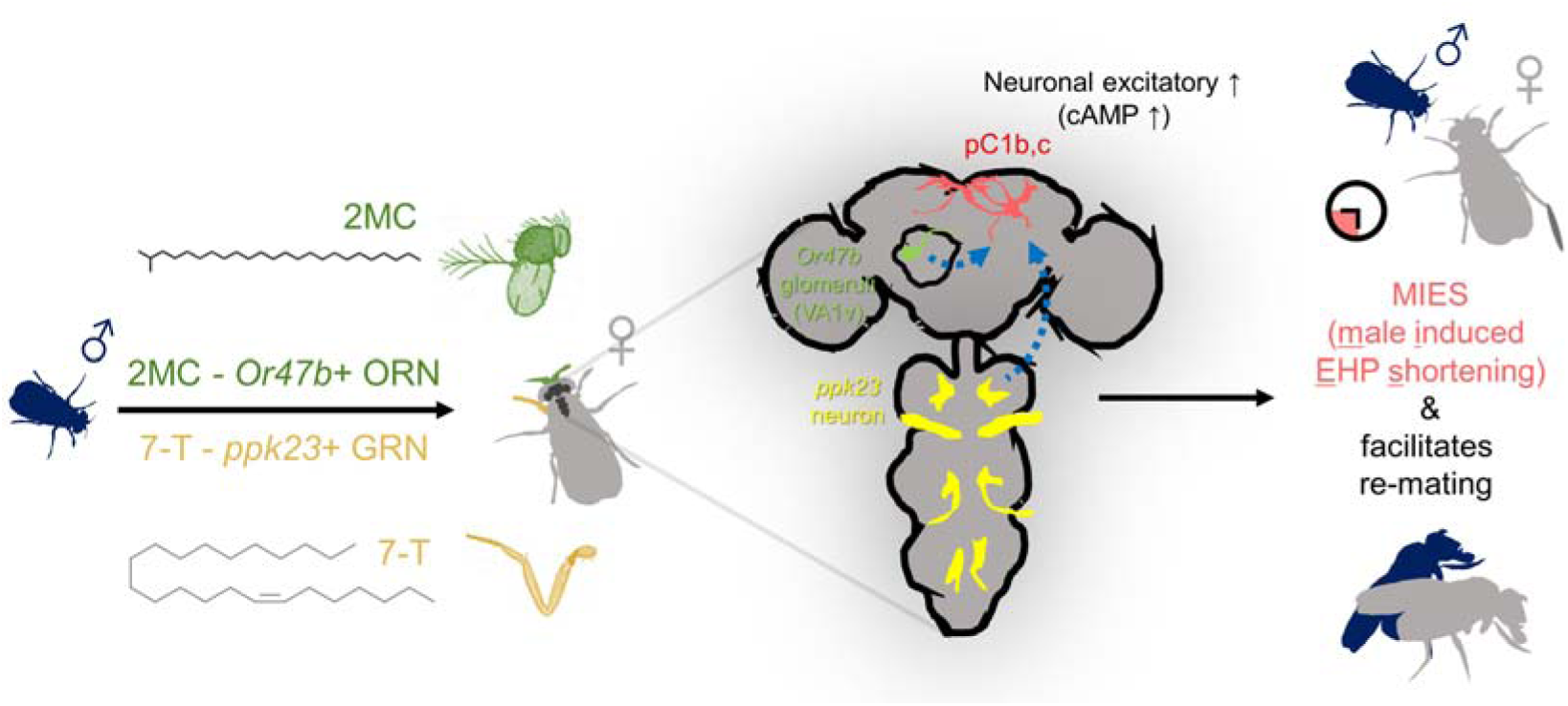
The presence of male odorants, which reflect changes in the social-sexual context, stimulates newly mated females to remove the male ejaculate and engage in subsequent re-mating. Following the initial mating, a female that encounters a new courting male removes the male ejaculate after a shorter EHP than those that do not encounter new male partners. This phenomenon, referred to as MIES in this study, is followed by a second mating with the new partner. The production of MIES depends on the functions of the Or47b+ olfactory and ppk23+ gustatory neurons, which are activated by 2MC and 7-T, respectively. These odorants increase cAMP levels in pC1b, c neurons, enhancing their responsiveness to male courtship cues and increasing mating receptivity. Consequently, 2MC and 7-T promote a second mating with a faster removal of the male ejaculate or mating plug.

## Discussion

Males employ a diverse range of strategies to enhance their reproductive fitness. One such strategy involves the formation ‘f a ’mating’plug’, a mechanism that prevents females from engaging in further mating, thereby increasing fertilization success rates (65–67). As a means of intra-sexual competition, rival males often promote the removal or precocious expulsion of the mating plug. The evolution of this strategy is driven by intersexual interactions with polyandrous females, who often remove the mating plug to engage in additional mating with males of superior traits or higher social status than their previous partners (37, 68). In the dunnock *Prunella modularis*, a small European passerine bird, the male often engages in cloacal pecking of mated females, inducing the expulsion of the previous ’mate’s sperm and mating plug, thereby increasing their chances of successful mating (39). In this study, we discovered that in *D. melanogaster*, freshly mated females exhibit an earlier removal of the mating plugs or a shorter EHP when kept with actively courting males. This behavior is primarily induced by the stimulation of females via male sex pheromones. In addition, our study has revealed that the neural circuit that processes male courtship cues and controls mating decisions plays an important role in regulating this behavior. This fly circuit has recently been proposed to be homologous to VMHvl in the mouse brain (45, 46). By delving into the molecular and neuronal mechanisms underlying MIES, our study provides valuable insights into the broader aspect of behaviors induced by changes in the social sexual context.

Our findings highlight the involvement of the Or47b receptor and Or47b ORNs in MIES. These OR and ORNs have been implicated in a range of social and sexual behaviors in both male and female fruit flies (12, 26–31). Methyl laurate and *trans*-palmitoleic acid are odorant ligands for Or47b that account for many of these functions particularly in males (30, 31). In this study, we provide compelling evidence that 2MC induces cAMP elevation in pC1 neurons and EHP shortening via both the Or47b receptor and Or47b ORNs, suggesting that 2MC functions as an odorant ligand for Or47b. Notably, gas chromatography–mass spectrometry (GC-MS) analysis of cuticular hydrocarbons from 4-day-old wild-type *D. melanogaster* revealed the presence of 2MC exclusively in males (30). Surprisingly, however, unlike 2MC, neither methyl laurate nor *trans*-palmitoleic acid affected EHP. The reason for this paradoxical result remains unclear. A plausible interpretation is that the EHP shortening induced by 2MC may require not only Or47b but also other as yet unidentified ORs. With the establishment of a behavioral and cellular assessment of 2MC activity, the search for additional odorant receptors responsive to 2MC is now feasible. Another important avenue for further research is whether 2MC can also elicit behaviors previously associated with methyl laurate or *trans*-palmitoleic acid, such as promoting male copulation and courtship (30, 31).

We observed that both 2MC and 7-T exhibit both cellular and behavioral activity within a specific concentration range (Fig. S4, S5, S7). This observation is of particular interest, given the multitude of environmental and biological factors that influence the levels of 2MC and 7-T, potentially affecting the capacity of males to induce MIES. For instance, exposure to low temperatures during development has been linked to increased production of both 2MC and 7-T (69). Similarly, mutation of the desiccation stress gene CG9186, which encodes a protein associated with lipid droplets, has been found to impact 2MC levels (70). Furthermore, 2MC levels rise with age in males (71). Thus, we propose that levels of 2MC, and possibly 7-T, may serve as indicators of male age and resilience to environmental stress in a complex manner.

In mated females, treatment with 2MC or 7-T increases cAMP levels in pC1b, c neurons but not in pC1a neurons. In contrast, pC1a neurons in virgin females are fully responsive to both male pheromones, showing increases in cAMP levels that are similar to those of pC1b,c neurons (Fig. S8). The absence of cAMP levels in pC1a neurons in mated females likely results from the mating signal (i.e., sex peptide) silencing pC1a neurons. Connectome and electrophysiology data support this interpretation, as SAG neurons, which relay sex peptide signals, have the strongest synaptic connection with the pC1a among five pC1 subtypes (49). However, the activity of SAG neurons may also influence pC1c neurons, as they also have substantial synaptic connections with pC1c neurons, as seen in the hemibrain connectome dataset (72). Future studies are needed to understand the role of SAG neurons in the regulation of EHP.

We found that increased cAMP levels cause pC1b, c neurons in mated females, which are typically unresponsive to male courtship cues like cVA and pulse song, to become responsive and exhibit strong Ca^2+^ transients. Since pC1b, c neurons play a role in generating sexual drive and increasing female receptivity to male courtship, the 2MC- or 7-T-induced increases in cAMP are likely to control the removal of the mating plug and the engagement of mated females in further mating (Fig. 7). This hypothesis aligns well with the previous report that mating reduces the sensitivity of Or47b ORNs, which we found to be responsive to 2MC, leading to an increased preference for pheromone-rich males after mating (29).

Moreover, the finding that 2MC and 7-T induce cAMP levels in pC1b, c neurons in virgin females suggests that virgin females may also use 2MC and 7-T as odorant cues to assess male quality during their first mating. Indeed, females seem to evaluate male quality by the amount of 7-T, as increased 7-T promotes mating receptivity and shortens mating latency (20).

Physiological factors like the nutritional status of females prior to mating and the nutritional status of their mates have been shown to influence EHP (73), and therefore potentially MIES. Hence, it is highly probable that MIES is regulated by additional central neurons such as Dh44-PI neurons that regulate these processes (40). However, it remains unclear whether and how Dh44-PI neurons and pC1 neurons interact to modulate EHP and MIES. The observation that double knockdown of Dh44R1 and Dh44R2 has only a marginal effect on MIES suggests that Dh44-PI neurons may also function independently of pC1 neurons, raising the possibility that multiple independent central circuits may contribute to the production of MIES.

Our initial screening of ORNs responsible for MIES revealed the involvement of Or47b ORNs, as well as several other ORNs. In addition to 2MC, which acts through Or47b-expressing ORNs, our findings indicate that 7-T and *ppk23* neurons, which are capable of detecting 7-T, also play a role in MIES induction. In *D. melanogaster* and other related species, food odors typically serve as volatile long-range signals that attract both males and females (74, 75), suggesting that specific food odors may also influence EHP (42). The involvement of multiple ORNs in the regulation of EHP predicts that pC1 neurons may process multiple odorants, not limited to those associated with mating behavior, including food odors. Future studies will explore the full spectrum of odorants processed by pC1 neurons in the regulation of EHP.

In conclusion, we have identified a circuit that, via the detection of a novel male pheromone, potentially signals male quality and governs the female’s decision to remove the mating plug of her last mate and mate again.

## Materials and Methods

### Fly care

Flies were cultured on a standard medium composed of dextrose, corn meal, and yeast, at room temperature on a 12hr : 12hr light:dark cycle (40, 73). Behavioral assays were performed at 25 °C, except for the thermogenetic activation experiment with dTRPA1. Virgin males and females were collected immediately after eclosion. Males were aged individually for 4-6 days, while females were aged in groups of 15–20. For EHP and mating assays, females were aged for 3-4 days. Assays were performed at Zeitgeber time (ZT) 3:00–11:00 and were repeated on at least three separate days.

### Fly stocks

The following stocks are from the Bloomington *Drosophila* Stock Center (BDSC), the Vienna *Drosophila* Resource Center (VDRC): *Canton S (CS)* (RRID: BDSC_64349), *w^1118^* (VDRC #60000), *R71G01* (pC1-Gal4) (RRID: BDSC_39599), *Orco1* (RRID: BDSC_23129), *Or13a-Gal4* (RRID: BDSC_9946), *Or19a-Gal4* (RRID: BDSC_9948), *Or23a-Gal4* (RRID: BDSC_9955), *Or43a-Gal4* (RRID: BDSC_9974), *Or47b-Gal4* (RRID: BDSC_9983), *Or47b-Gal4* (RRID: BDSC_9984), *Or65a-Gal4* (RRID: BDSC_9993), *Or65b-Gal4* (RRID: BDSC_23901), *Or65c-Gal4* (RRID: BDSC_23903), *Or67d-Gal4* (RRID: BDSC_9998), *Or83c-Gal4* (RRID: BDSC_23131), *Or88a-Gal4* (RRID: BDSC_23137), *UAS-Or47b* (RRID: BDSC_76045), *Or47b2/2* (RRID: BDSC_51306), *Or47b3/3* (RRID: BDSC_51307), *UAS-TNT active* (RRID: BDSC_28837), *UAS-TNT inactive* (RRID: BDSC_28839), *UAS-dTRPA1* (RRID: BDSC_26263), *UAS-CsChrimson* (RRID: BDSC_55135), *uAS-GCaMP6m* (RRID: BDSC_42748), *R52G04-AD* (RRID: BDSC_71085), *SAG-Gal4* (VT50405) (RRID:Flybase_FBst0489354, VDRC #200652), *UAS-Dh44R1-RNAi* (RRID:Flybase_FBst0482273, VDRC #110708), *UAS-Dh44R2-RNAi* (RRID:Flybase_FBst0465025, VDRC #43314), *UAS-Dicer2* (VDRC #60007). The following stocks were previously reported: *PromE*(*800*)*-Gal4* (59), *UAS-FLP, CRE-F-luc* (76), *LexAop-FLP* (77), *UAS-CsChrimson* (78), *UAS-GtACR1* (79), *UAS-PhotoAC* (PACα) (80), pC1-A (48), pC1-S (48), *Dh44-pC1-Gal4* (47), *ppk23*-*Gal4*, *ppk23*-, *ppk28*-, *ppk29*-(62), and *Orco-Gal4*, *UAS-EGFP-Orco* (81). *pC1a-split-Gal4* is generated by combining *R52G04-AD* (RRID: BDSC_71085) and *dsx-DBD* (49). *Drosophila* species other than *D. melanogaster* are obtained from the EHIME-Fly *Drosophila* Stock Center and the KYORIN-Fly *Drosophila* species Stock Center. To enhance knock-down efficiency, RNAi experiments were performed using flies carrying *UAS-Dicer2* (VDRC #60007).

### Chemical information

All *trans*-retinal (Cat# R2500), methyl laurate (Cat# W271500), and Triton™ X-100 (Cat# X100) were obtained from Sigma-Aldrich (St. Louis, MO, USA). The following chemicals were obtained from the Cayman Chemical (Ann Arbor, MI, USA): 7(Z)-Tricosene (CAS No. 52078-42-9, Cat# 9000313), 7(Z)-Pentacosene (CAS No. 63623-49-4, Cat# 9000530), *trans*-palmitoleic acid (CAS No. 10030-73-6, Cat# 9001798), 11-*cis*-vaccenyl acetate (cVA) dissolved in EtOH (CAS No. 6186-98-7, Cat# 10010101). 2-methyltetracosane (>98% purity) was custom-synthesized by KIP (Daejeon, Korea). Ethanol is used as a vehicle for 7-T, cVA, *trans*-palmitoleic acid and methyl laurate, while hexane is used as a vehicle for 2-methyltetracosane.

### Behavior assays

For mating behavior assays, we followed the procedures described previously (82). Individual virgin females and naive *CS* males were paired in 10 mm diameter chambers and were recorded using a digital camcorder (SONY, HDR-CX405 or Xiaomi, Redmi Note 10) for either 30 minutes or 1 hour for the mating assay and 6 hours for the re-mating assay. In the re-mating assay, females that completed their initial mating within 30 minutes were subsequently paired with naive *CS* males.

To measure EHP, defined as the time elapsed between the end of copulation and sperm ejection, we used the following procedure: Virgin females were individually mated with *CS* males in 10 mm diameter chambers. Following copulation, females were transferred to new chambers, either with or without a *CS* male or pheromone presentation, and their behavior was recorded using a digital camcorder (SONY, HDR-CX405). Typically, females that completed copulation within 30 minutes were used for analysis. The sperm ejection scene, in which the female expels a white sac containing sperm and the mating plug through the vulva, was directly observed by eye in the recorded video footage. For pheromone presentation, females were individually housed in 10 mm diameter chambers containing a piece of Whatman filter paper (2 mm x 2 mm) treated with 0.5 μl of the pheromone solution and air dried for 1 minute. For thermogenetic activation experiments, females were incubated at the indicated temperatures immediately after the end of copulation. For light activation experiments, a custom-made light activation setup was used with a ring of 104 multi-channel LED lights (NeoPixel, Cat# WS2812; red light, 620-625 nm, 390-420 mcd; green light, 522-525 nm, 660-720 mcd; blue light, 465-467 nm, 180-200 mcd). Females were individually placed in 10 mm diameter chambers, and the chamber was illuminated with light at an intensity of 1100 lux across the chamber during the assay, as measured by an HS1010 light meter. Flies used in these experiments were prepared by culturing them immediately after eclosion in food containing vehicle (EtOH) or 1 mM all *trans*-retinal (ATR). They were kept in complete darkness for 3-4 days until the assay was conducted. To prevent the accumulation of residual pheromones, all behavioral chambers were cleaned with 70% water/ethanol or acetone before and after the experiment.

### Calcium imaging

We followed the procedures described previously (47, 83). Following copulation, freshly mated female flies were temporarily immobilized using ice anesthesia, and their heads were attached to a custom-made thin metal plate with a 1-mm diameter hole using photo-curable UV glue (ThreeBond, A16A01). An opening in the fly’s head was created using a syringe needle under saline (108□mM NaCl, 5□mM KCl, 2□mM CaCl_2_, 8.2□mM MgCl_2_, 4□mM NaHCO_3_, 1□mM NaH_2_PO_4_, 5□mM trehalose, 10□mM sucrose, 5□mM HEPES pH 7.5). Imaging was performed with a Zeiss Axio Examiner A1 microscope equipped with an electron-multiplying CCD camera (Andor Technology, Luca^EM^ R 604M) and an LED light source (CoolLED, Precis Excite). Metamorph software (Molecular Devices, RRID:SCR_002368) was used for image analysis. The Syntech Stimulus Controller (Type CS-55) was used to deliver the male pheromone using an airflow. 2 μl of pheromone solution was applied to a piece of Whatman filter paper (2 mm x 1 mm), which was then inserted into a glass Pasteur pipette after solvent evaporation.

### Luciferase assay

We followed the procedures described previously (47, 76). For the assay, 3-day-old virgin females or freshly mated females were used. A group of three fly heads, kept at -80°C, was homogenized using cold homogenization buffer (15 mM HEPES, 10 mM KCl, 5 mM MgCl_2_, 0.1 mM EDTA, 0.5 mM EGTA).

Luciferase activity was measured using beetle luciferin potassium salt (Promega, Cat# E1603) and a microplate luminometer (Berthold technologies, Centro XS^3^ LB 960), following the manufacturer’s instructions. For pheromone presentation, flies were placed in 10 mm diameter chambers containing a piece of Whatman filter paper (4 mm x 6 mm) treated with 1 μl of the pheromone solution and air dried for 1 minute.

### Immunohistochemistry

3–5-day-old virgin female flies were dissected in phosphate buffered saline (PBS) and fixed in 4% paraformaldehyde in PBS for 30 minutes at room temperature. After fixation, the brains were thoroughly washed in PBST (0.1% Triton™ X-100 in PBS) and then blocked with 5% normal goat serum in PBST. After blocking, brains were incubated with primary antibody in PBST for 48 hours at 4°C, washed with PBST, and then incubated with secondary antibody in PBST for 24 hours at 4°C. The samples were washed three times with PBST and once with PBS before mounting in Vectashield (Vector Laboratories, Cat# H-1000). Antibodies used were rabbit anti-GFP (1:1000; Thermo Fisher Scientific, Cat# A-11122, RRID:AB_221569), mouse anti-nc82 (1:50; Developmental Studies Hybridoma Bank, Cat# Nc82; RRID: AB_2314866), Alexa 488-conjugated goat anti-rabbit (1:1000; Thermo Fisher Scientific, Cat# A-11008, RRID:AB_143165), Alexa 568-conjugated goat anti-mouse (1:1000; Thermo Fisher Scientific, Cat# A-11004, RRID:AB_2534072). Brain images were acquired with a Zeiss LSM 700/Axiovert 200M (Zeiss) and processed with Fiji (https://imagej.net/software/fiji/downloads, RRID:SCR_002285)

### Color depth MIP based anatomical analysis

A stack of confocal images of *pC1a-split-Gal4*>*UAS-myr-EGFP* adult female brains stained with anti-GFP and anti-nc82 was used. Images were registered to the JRC2018 unisex brain template (84) using the Computational Morphometry Toolkit (CMTK, https://github.com/jefferis/fiji-cmtk-gui). Color depth MIP masks of *pC1a-split-Gal4* neurons and pC1a (ID, 5813046951) in Hemibrain (72) (Fig. S6C) were generated using the ColorMIP_Mask_Search plugin (85) for Fiji (https://github.com/JaneliaSciComp/ColorMIP_Mask_Search) and NeuronBridge (86) (https://neuronbridge.janelia.org/). Similarity score and rank were calculated using NeuronBridge.

### Statistical analysis

Statistical analysis was conducted using GraphPad Prism 9 (Graphpad, RRID:SCR_002798), with specific details of each statistical method provided in the figure legends.

## Acknowledgments

We thank S. Kang, J-H. Yoon, and B. Lee for excellent technical assistance and the GIST Advanced Institute of Instrumental Analysis (GAIA) for the confocal microscopy analysis. Fly stocks were obtained from the Bloomington *Drosophila* Stock Center (NIH P40OD018537), the Vienna *Drosophila* Resource Center (VDRC), the Kyoto Stock Center, the EHIME-Fly *Drosophila* species stock center, the KYORIN-Fly *Drosophila* species stock center, and the Korea *Drosophila* Resource Center (NRF-2022M3H9A1085169). This work was supported by National Research Foundation of Korea grants NRF-2022R1A2C3008091 (Y-J.K.), 2022M3E5E8081194 (Y-J.K.), NRF-2019R1A4A1029724 (Y-J.K.), 2017R1A6A3A11027866 (D-H.K.), NRF-2021R1I1A1A01060304 (D-H.K.), GIST Research Institute (GRI) GIST-MIT research collaboration grant funded by GIST in 2023 (Y-J.K.), 2022, 2023 AI-based GIST Research Scientist Project (D-H.K.).

## Author contributions

Conceptualization: M.Y., Y-J.K., Methodology: K-M.L., M.K., B.S.H., Investigation: M.Y., D-H.K., T.S.H., E.P., Visualization: M.Y., Supervision: Y-J.K., Writing—original draft: M.Y., Y-J.K., Writing—review & editing: M.Y., M.K., B.S.H., Y-J.K.

## Competing interests

The authors declare that they have no competing interest.

## Notes

### Competing Interest Statement

The authors have declared no competing interest.

### Summary of Updates

In the revised manuscript, we made the following changes. 1.We provided new evidence that Or47b is required for 2MC-induced cAMP increase in pC1 neurons, but not for 7T-induced increase. This observation supports that Or47b functions as a receptor for 2MC. 2.We showed the expression of the pC1a-split-Gal4 transgene in the brain and VNC. 3.We prepared a new figure summarizing the results and explaining the current working model.4.We clarified the comparisons made in the multiple comparison analysis and specified the tests used in several figures.5.We corrected typographical errors and made some changes to improve readability.

