## Supplemental Information for "Male cuticular pheromones stimulate removal of the mating plug and promote re-mating through pC1 neurons in *Drosophila* females"

**This PDF file includes:**

Figures S1 to S9
Table S1

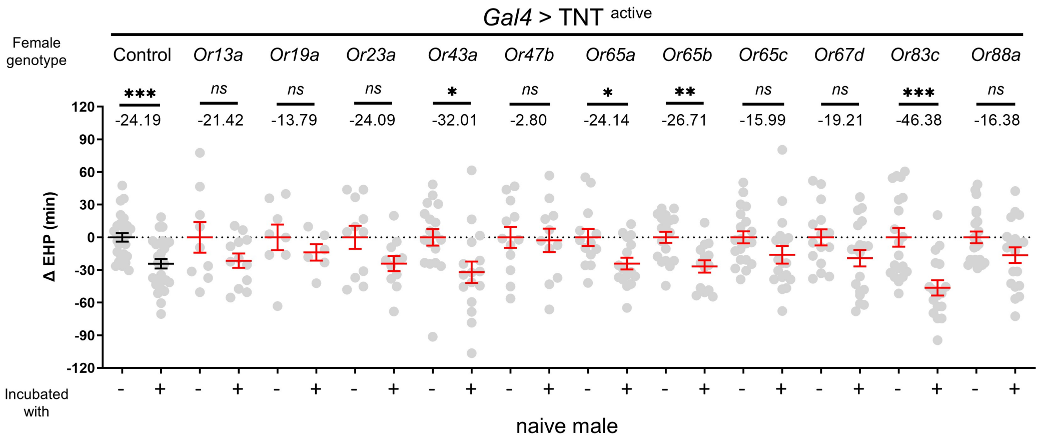


**Fig. S1. The identification of trichoid and intermediate sensilla ORNs that are necessary for the production of MIES**

Δ EHP of females of the indicated genotypes, incubated with or without naive males immediately after mating. The female genotypes are as follows from left to right: *+>*$TNT^{active}$, *Or13a>*$TNT^{active}$, *Or19a>*$TNT^{active}$, *Or23a>*$TNT^{active}$, *Or43a>*$TNT^{active}$, *Or47b>*$TNT^{active}$, *Or65a>*$TNT^{active}$, *Or65b>*$TNT^{active}$, *Or65c>*$TNT^{active}$, *Or67d >*$TNT^{active}$, *Or83c >*$TNT^{active}$, *Or88a>*$TNT^{active}$.

Mann-Whitney Test (n.s. $p > 0.05$; *$p < 0.05$; **$p < 0.01$; ***$p < 0.001$). Gray circles indicate the ΔEHP of individual females, and the mean ± SEM of data is presented. The ΔEHP is calculated by subtracting the mean of the reference EHP of females kept alone after mating ('-') from the EHP of individual females in comparison. Numbers below the horizontal bar represent the mean of the EHP differences between the indicated treatments.

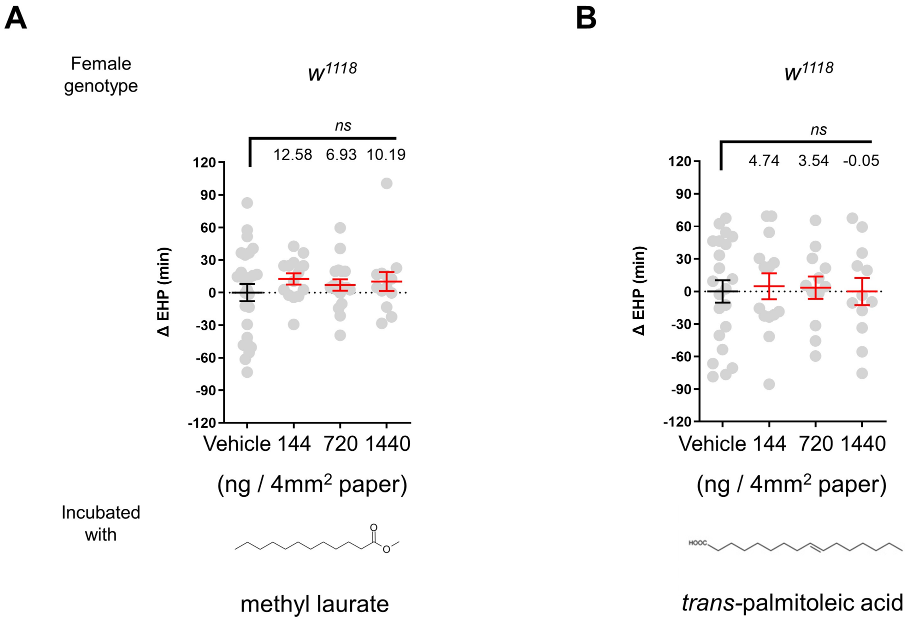


**Fig. S2. Known odorant ligands for *Or47b*, methyl laurate and *trans*-palmitoleic acid, were unable to induce EHP shortening**

ΔEHP of $w^{1118}$ females incubated with a piece of filter paper perfumed with solvent vehicle or with the indicated amounts of two known *Or47b* odorant ligands, methyl laurate (A) and *trans*-palmitoleic acid (B) immediately after mating. Mann-Whitney Test (n.s. $p > 0.05$). The ΔEHP is calculated by subtracting the mean of the reference EHP of females incubated with vehicle-perfumed paper (the leftmost column) from the EHP of individual females in comparison. Gray circles indicate the ΔEHP of individual females, and the mean ± SEM of data is presented. Numbers below the horizontal bar represent the mean of the EHP differences between vehicle and odorant treatments.

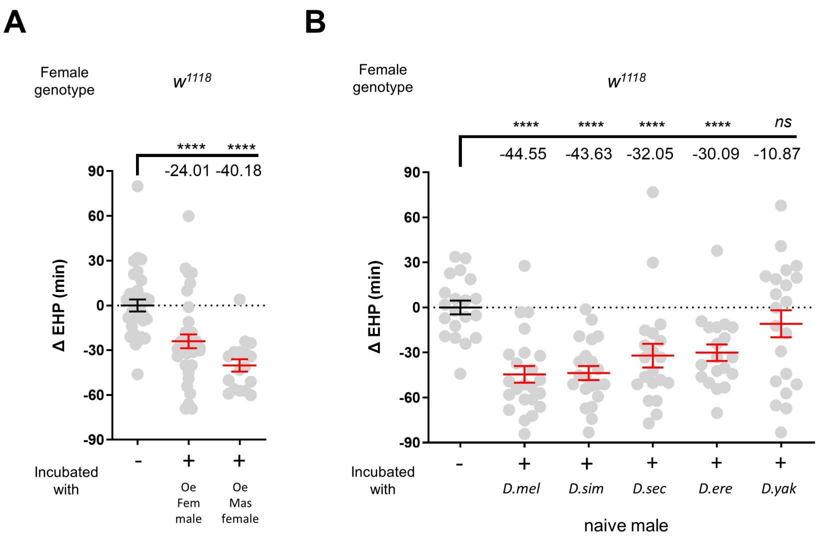


**Fig. S3. EHP shortening is induced by males with feminized oenocytes, females with masculinized oenocytes, and males of other closely related *Drosophila* species**

**A**, ΔEHP of $w^{1118}$ females incubated with males with feminized oenocytes (Oe Fem male; *PromE(800)-Gal4/UAS-Tra*) or virgin females with masculinized oenocytes (Oe Mas Female; *PromE(800)-Gal4/UAS-Tra-RNAi*).

**B**, Δ EHP of $w^{1118}$ females incubated with naive males of the indicated *Drosophila* species. *D. mel* (*D. melanogaster*), *D. sim* (*D. simulans*), *D. sec* (*D. sechellia*), *D. ere* (*D. erecta*), *D. yak* (*D. yakuba*). Mann-Whitney Test (n.s. $p > 0.05$; ****$p < 0.0001$). The ΔEHP is calculated by subtracting the mean of the reference EHP of females kept alone after mating (the leftmost column) from the EHP of individual females in comparison. Gray circles indicate the ΔEHP of individual females, and the mean ± SEM of data is presented. Numbers below the horizontal bar represent the mean EHP differences between the indicated treatments.

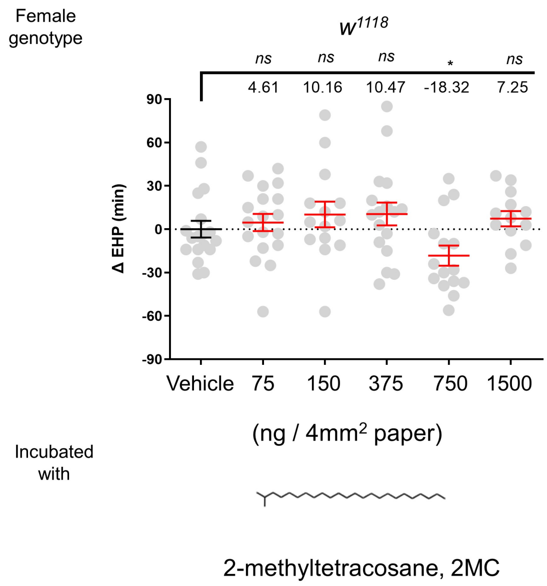


**Fig. S4. 2MC shortens EHP at a specific concentration**

ΔEHP of *w*$^{1118}$ females incubated with a piece of filter paper perfumed with solvent vehicle or the indicated amounts of 2MC. Mann-Whitney Test (n.s. $p > 0.05$; *$p < 0.05$). The ΔEHP is calculated by subtracting the mean of the reference EHP of females incubated with vehicle-perfumed paper (the leftmost column) from the EHP of individual females in comparison. Gray circles indicate the ΔEHP of individual females, and the mean ± SEM of data is presented. Numbers below the horizontal bar represent the mean of the EHP differences between vehicle and odorant treatments.

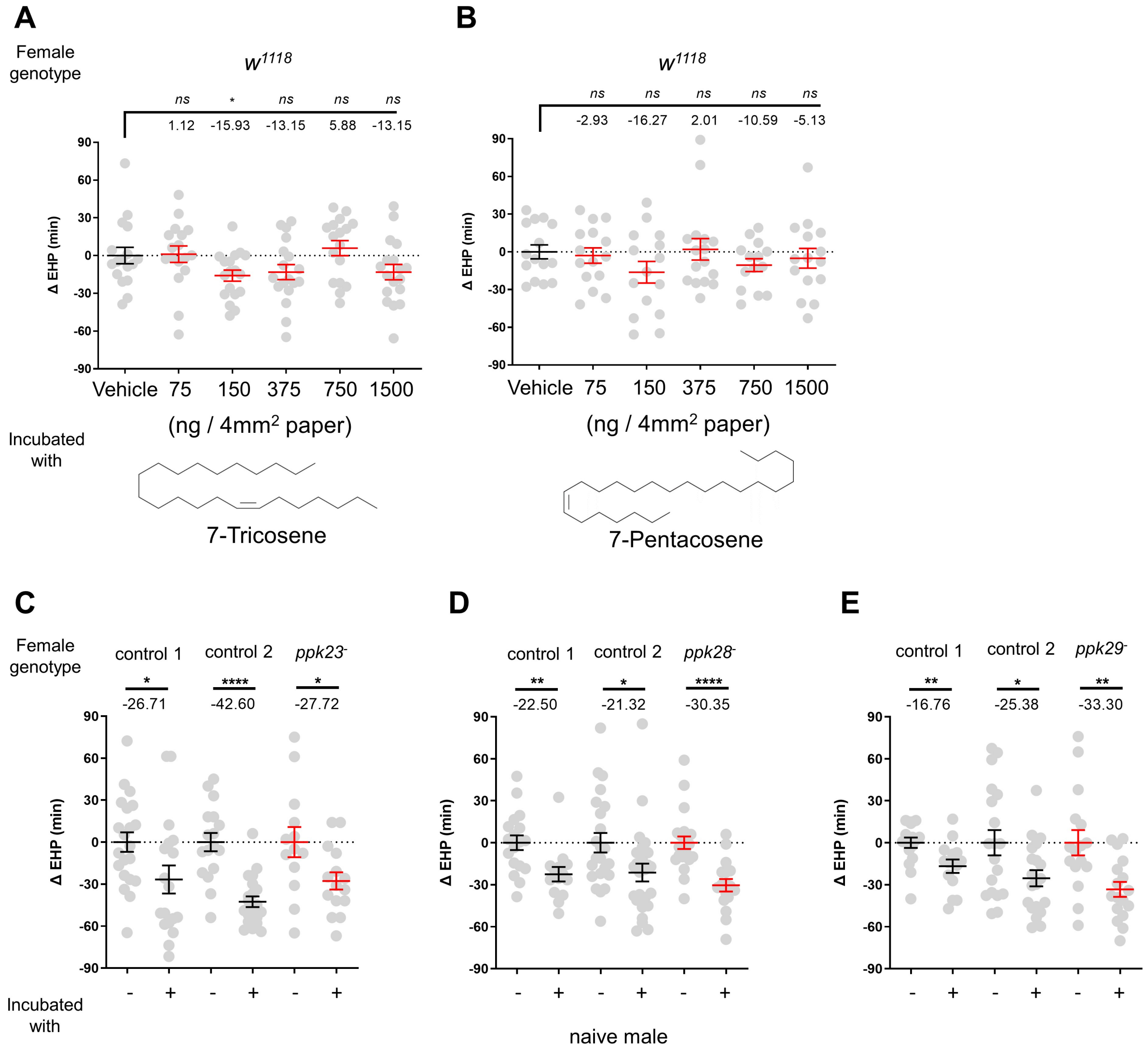


**Fig. S5. 7-T induces EHP shortening at physiological concentrations, but DEG/ENaC channels expressed in *ppk23* neurons are not required for MIES**

**A-B**, ΔEHP of $w^{1118}$ females incubated with a piece of filter paper perfumed with solvent vehicle or the indicated amounts of 7-T (A), or 7-Pentacosene (B) after mating. Incubation with a specific concentration of 7-T significantly shorten EHP, but 7-Pentacosene does not. Unpaired *t*-Test (n.s. $p > 0.05$; *$p < 0.05$).
**C-E**, ΔEHP of females of the indicated genotypes, incubated with or without naive males after mating. The female genotypes are as follows from left to right: (C) control 1 ($w^{1118}$), control 2 ($ppk23^{-}/+$), and $ppk23^{-}$ ($ppk23^{-}/ppk23^{-}$); (D) control 1 ($w^{1118}$), control 2 ($ppk28^{-}/+$), and $ppk28^{-}$ ($ppk28^{-}/ppk28^{-}$); (E) control 1 ($w^{1118}$), control 2 ($ppk29^{-}/+$), and $ppk29^{-}$ ($ppk29^{-}/ppk29^{-}$). Mann-Whitney Test (n.s. $p > 0.05$; *$p < 0.05$; **$p < 0.01$; ****$p < 0.0001$). The ΔEHP is calculated by subtracting the mean of the reference EHP of females incubated with vehicle-perfumed paper (the leftmost column in A, B) or kept alone after mating ('-' in C-E) from the EHP of individual females in comparison. Gray circles indicate the ΔEHP of individual females, and the mean ± SEM of data is presented. Numbers below the horizontal bar represent the mean of the EHP differences between the indicated treatments.

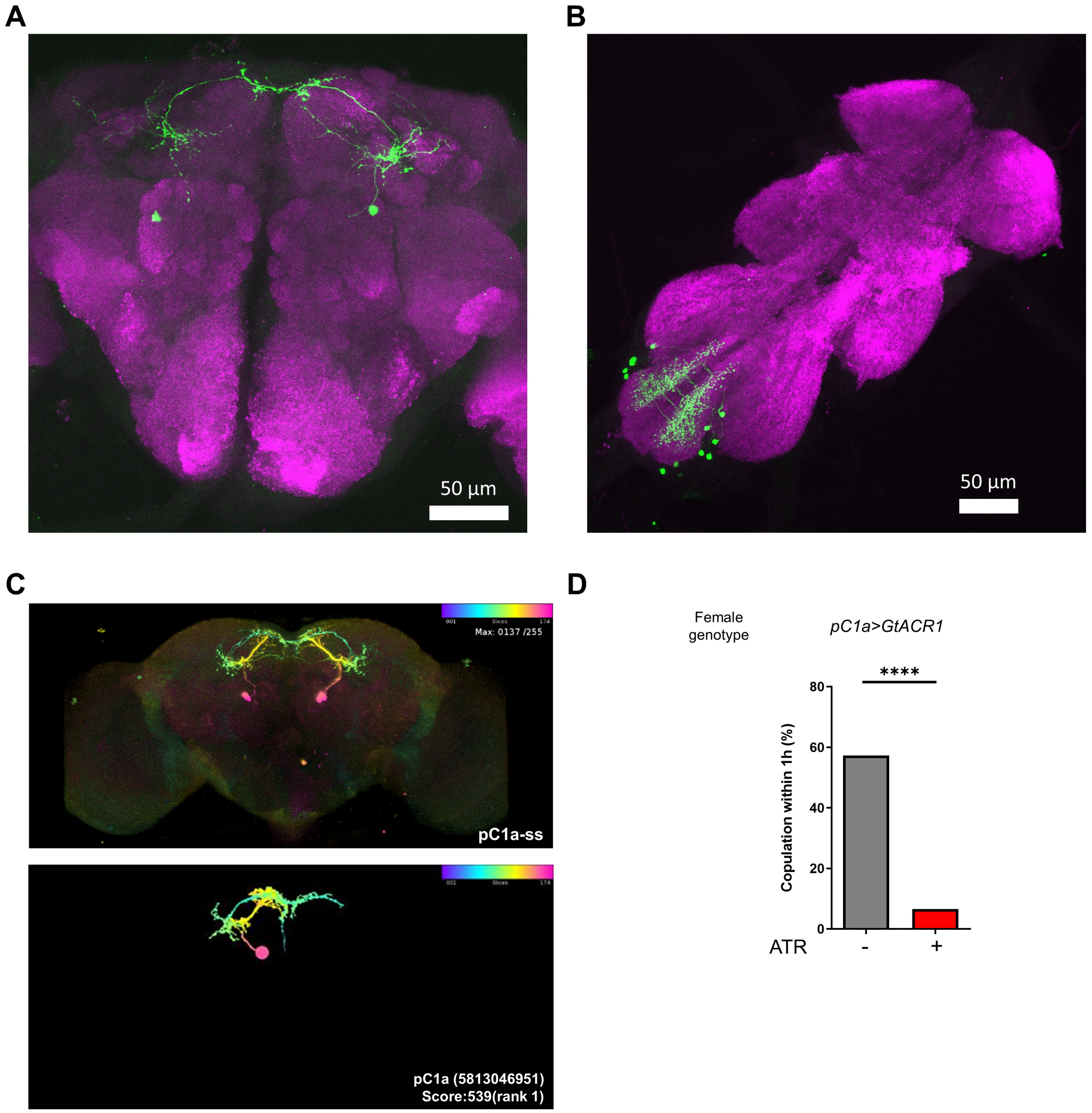


**Fig. S6. Characterization of *pC1a-split-Gal4***

**A, B,** Z-projection confocal images of the brain (A) and VNC (B) of a female carrying *pC1a-split-GAL4* and *UAS-myrEGFP*, stained with anti-EGFP (green) and anti-nc82 (magenta). Scale bars, 50 µm. In the brain, only the pC1a cells are labeled, but in the VNC, several cells are labeled in the abdominal ganglion.

**C**, An anatomical comparison between *pC1a-split-GAL4* neurons (above; pC1a-ss) in the brain and a pC1a neuron (below; neuprint body ID, 5813046951). The panel above shows the maximum intensity projection image (MIP) of an aligned confocal image of the brain from a female carrying *pC1a-split-GAL4* and *UAS-myrEGFP* stained with anti-EGFP and anti-nc82.

**D**, Mating frequencies of *pC1a>GtACR1* (*pC1a-split-Gal4/UAS-GtACR1*) females during optogenetic silencing, scored as the percentage of females that copulate within 1 hour. Females were cultured on food with or without all *trans*-retinal (ATR) prior to the mating assay. The optogenetic silencing of pC1a neurons with GtACR1 was observed to suppress mating receptivity almost completely. Chi-square test (****$p < 0.0001$).

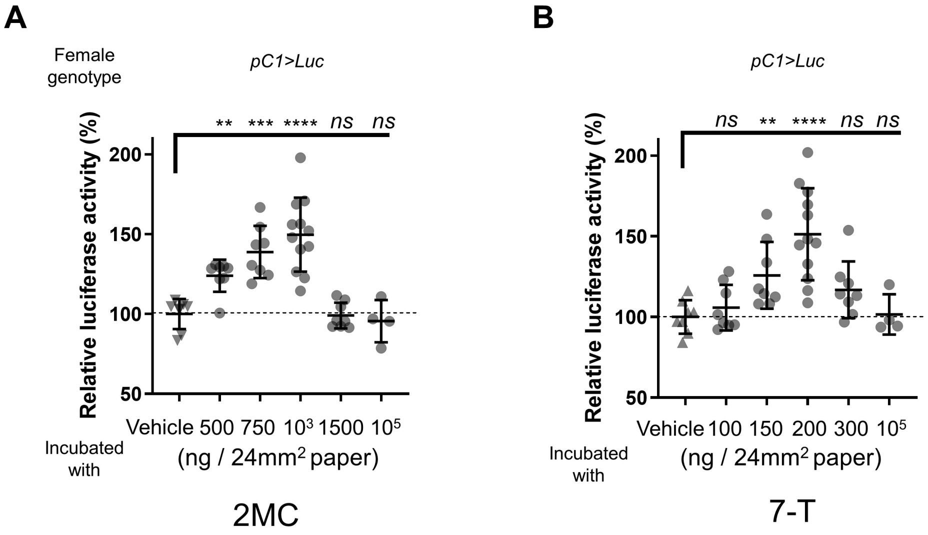


**Fig. S7. Incubation with 2MC or 7-T increases cAMP levels in pC1 neurons**
The relative *CRE-luciferase* reporter activity of pC1 neurons in females incubated with a piece of filter paper perfumed with the indicated amounts of 2MC (A) and 7-T (B). It is noteworthy that the concentration range within which 2MC or 7-T increases cAMP levels in pC1 neurons is narrow. To calculate the relative luciferase activity, the average luminescence unit values of the female incubated with the vehicle are set to 100%. Gray circles indicate the relative luciferase activity (%) of individual females, and the mean ± SEM of data is presented. Mann-Whitney Test (n.s. $p > 0.05$; **$p < 0.01$; ***$p < 0.001$; ****$p < 0.0001$).

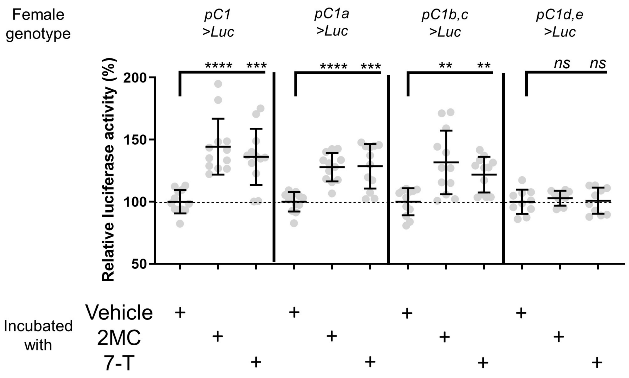


**Fig. S8. Incubation with 2MC or 7-T increases cAMP levels in pC1a as well as pC1b, c neurons in virgin females**

The relative *CRE-luciferase* reporter activity of pC1 neurons in virgin females of the indicated genotypes, incubated with a piece of filter paper perfumed with the indicated odorants. To calculate the relative luciferase activity, the average luminescence unit values of the female incubated with the vehicle are set to 100%. Mann-Whitney Test (n.s. $p > 0.05$; **$p < 0.01$; ***$p < 0.001$; ****$p < 0.0001$). Gray circles indicate the relative luciferase activity (%) of individual females, and the mean ± SEM of data is presented.

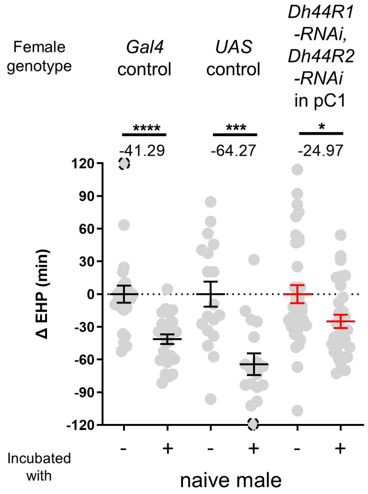


**Fig. S9. The knockdown of Dh44R1 and Dh44R2 in pC1 neurons has a limited impact on MIES**
ΔEHP of females of the indicated genotypes, incubated with or without naive males immediately after mating. The female genotypes are as follows from left to right: *Gal4* control (*UAS-Dcr2/+; GMR71G01-Gal4/+*), *UAS* control (*UAS-Dh44R1-RNAi/+; UAS-Dh44R2-RNAi/+*), *Dh44R1-RNAi, Dh44R2-RNAi* in pC1 (*UAS-Dcr2/+; GMR71G01-Gal4/Dh44R1-RNAi; Dh44R2-RNAi/+*). Mann-Whitney Test (*$p$ <0.05; ***$p$ < 0.001; ****$p$ < 0.0001). The ΔEHP is calculated by subtracting the mean of the reference EHP of females kept alone after mating ('-') from the EHP of individual females in comparison. Gray circles indicate the ΔEHP of individual females, and the mean ± SEM of data is presented. The gray circles with dashed borders indicate ΔEHP values that exceed the axis limits (>120 or <-120 minutes). Numbers below the horizontal bar represent the mean of the EHP differences between the indicated treatments.

| Figure | Genotype of | | | N numbers |
|---|---|---|---|---|
| | Tested females | Mating partner | Incubation partner | From left to right |
| Fig. 1 | | | | |
| Fig. 1B | *w[1118]* | *Canton-S* | *Canton-S* | 59, 69 |
| Fig. 1C | *w[1118]* | *Canton-S* | *Canton-S* | 18, 15 |
| Fig. 1D | *w[1118]* | *Canton-S* | *Canton-S* | 12, 12 |
| Fig. 1E | *w[1118]* | *Canton-S* | *Canton-S* | 20, 18, 23 |
| Fig. 1F | *w[1118]; TI{w[+mW.hs]=TI}Orco[1]* | *Canton-S* | *Canton-S* | 55, 47 |
| Fig. 2 | | | | |
| Fig. 2A | *w[1118];Or47b-Gal4/UAS-TNTinactive(P{UAS-TeTxLC.(-)Q}A2)* | *Canton-S* | *Canton-S* | 14, 14 |
| | *w[1118];Or47b-Gal4/UAS-TNTactive (P{w[+mC]=UAS-TeTxLC.tnt}E2)* | *Canton-S* | *Canton-S* | 16, 18 |
| Fig. 2B | *w[1118];Or47b-Gal4/+* | *Canton-S* | *Canton-S* | 28, 31 |
| | *w[1118];;UAS-dTRPA1/+* | *Canton-S* | *Canton-S* | 21, 16 |
| | *w[1118];Or47b/+;UAS-dTRPA1/+* | *Canton-S* | *Canton-S* | 11, 15 |
| Fig. 2C | *w[1118]; TI{w[+mW.hs]=TI}Orco[1]* | *Canton-S* | *Canton-S* | 12, 12 |
| | *w[1118]; Or47b-Gal4>UAS-EGFP-Orco; Orco[1]/Orco[1]* | *Canton-S* | *Canton-S* | 13, 14 |
| Fig. 2D | *w[1118]; Or47b[2]/+* | *Canton-S* | *Canton-S* | 12, 15 |
| | *w[1118]; Or47b[3]/+* | *Canton-S* | *Canton-S* | 13, 14 |
| | *w[1118]; Or47b[2]/Or47b[3]* | *Canton-S* | *Canton-S* | 13, 12 |
| Fig. 2E | *w[1118]; Or47b[2]/Or47b[2]* | *Canton-S* | *Canton-S* | 14, 15 |
| | *w[1118]; Or47b-Gal4>P{w[+mC]=UAS-Or47b.MYC}2; Or47b[2]/Or47b[2]* | *Canton-S* | *Canton-S* | 11, 11 |
| Fig. 3 | | | | |
| Fig. 3A | *w[1118]* | *Canton-S* | | 13, 16 |
| Fig. 3B | *w[1118]; TI{w[+mW.hs]=TI}Orco[1]* | *Canton-S* | | 11, 12 |
| Fig. 3C | *w[1118]; TI{w[+mW.hs]=TI}Or47b[2]* | *Canton-S* | | 14, 17 |
| Fig. 3D | *w[1118];Or47b-Gal4/+;Orco[1]/Orco[1]* | *Canton-S* | | 22, 22 |
| | *w[1118];UAS-Orco/+; Orco[1]/Orco[1]* | *Canton-S* | | 15, 14 |
| | *w[1118];Or47b-Gal4/UAS-Orco;Orco[1]/Orco[1]* | *Canton-S* | | 18, 19 |
| Fig. 4 | | | | |
| Fig. 4A | *w[1118]* | *Canton-S* | *Canton-S* | 18, 22 |
| Fig. 4B | *w[1118]* | *Canton-S* | | 23, 31 |
| Fig. 4C | *w[1118]* | *Canton-S* | | 16, 17 |

| | | | | |
|---|---|---|---|---|
| Fig. 4D | w[1118];ppk23-Gal4/UAS-TNTinactive(P{UAS-TeTxLC.(-)Q}A2) | Canton-S | Canton-S | 18, 13 |
| | w[1118];ppk23-Gal4/UAS-TNTactive (P{w[+mC]=UAS-TeTxLC.tnt}E2) | Canton-S | Canton-S | 17, 17 |

| | | | | |
|---|---|---|---|---|
| Fig. 5 | | | | |
| Fig. 5A | w[1118];pC1(R71G01)-AD/+;Dsx-DBD/UAS-GtACR1 | Canton-S | | 22, 21 |
| Fig. 5B | w[1118];VT25602-AD/+;UAS-GtACR1/VT2064-DBD | Canton-S | | 18, 18 |
| Fig. 5C | w[1118];R52G04-AD/+;UAS-GtACR1/Dsx-DBD | Canton-S | | 15, 14 |
| Fig. 5D | w[1118];;Dh44-pC1 (Dsx-DBD, Dh44A-AD)-GAL4/UAS-GtACR1 | Canton-S | | 17, 20 |
| Fig. 5E | w[1118];UAS-FLP/+; GMR71G01-Gal4, CRE-F-Luc/+ | Canton-S | | 12, 12, 12 |
| | w[1118];R52G04-AD/+;UAS-FLP, CRE-F-Luc/Dsx-DBD | Canton-S | | 12, 12, 12 |
| | w[1118];;UAS-FLP, CRE-F-Luc/Dh44-pC1 (Dsx-DBD, Dh44A-AD)-GAL4 | Canton-S | | 16, 16, 16 |
| | w[1118];VT25602-AD/+;UAS-FLP, CRE-F-Luc/VT2064-DBD | Canton-S | | 12, 12, 12 |
| Fig. 5F | w[1118]; Or47b[2]/+; pC1-FLP, CRE-F-Luc | Canton-S | | 8, 8, 8 |
| | w[1118]; Or47b[2]/Or47b[2]; pC1-FLP, CRE-F-Luc | Canton-S | | 8, 8, 8 |
| | w[1118]; Or47b[3]/+; pC1-FLP, CRE-F-Luc | Canton-S | | 8, 8, 8 |
| | w[1118]; Or47b[3]/Or47b[3]; pC1-FLP, CRE-F-Luc | Canton-S | | 8, 8, 8 |

| | | | | |
|---|---|---|---|---|
| Fig. 6A | w[1118]; R52G04-AD/+;UAS-PhotoAC/Dsx-DBD | Canton-S | | 18, 22 |
| | w[1118];;UAS-PhotoAC/Dh44-pC1 (Dsx-DBD, Dh44A-AD)-GAL4 | Canton-S | | 22, 28 |
| | w[1118]; VT25602-AD/+;UAS-PhotoAC/VT2064-DBD | Canton-S | | 21, 20 |
| Fig. 6B | w[1118];UAS-GCaMP6m/+; pC1(GMR71G01)-GAL4/UAS-PhotoAC | Canton-S | | 9, 9, 9 |
| Fig. 6C | w[1118];;+/UAS-PhotoAC | Canton-S (1st, 2nd) | | 60 |
| | w[1118];;Dh44-pC1 (Dsx-DBD, Dh44A-AD)-GAL4/UAS-PhotoAC | Canton-S (1st, 2nd) | | 18 |

| | | | | |
|---|---|---|---|---|
| Fig. S1 | w[1118];+/P{w[+mC]=UAS-TeTxLC.tnt}E2 | Canton-S | Canton-S | 27, 27 |
| | w[1118];Or13a-Gal4/P{w[+mC]=UAS-TeTxLC.tnt}E2 | Canton-S | Canton-S | 9, 12 |
| | w[1118];+/P{w[+mC]=UAS-TeTxLC.tnt}E2;+/Or19a-Gal4 | Canton-S | Canton-S | 8, 6 |
| | w[1118];+/P{w[+mC]=UAS-TeTxLC.tnt}E2;+/Or23a-Gal4 | Canton-S | Canton-S | 11, 11 |
| | w[1118];+/P{w[+mC]=UAS-TeTxLC.tnt}E2;+/Or43a-Gal4 | Canton-S | Canton-S | 18, 16 |

| | | | | |
|---|---|---|---|---|
| | *w[1118];+/P{w[+mC]=UAS-TeTxLC.tnt}E2;+/Or47b-Gal4* | *Canton-S* | *Canton-S* | 12, 11 |
| | *w[1118];Or65a-Gal4/P{w[+mC]=UAS-TeTxLC.tnt}E2* | *Canton-S* | *Canton-S* | 13, 16 |
| | *w[1118];Or65b-Gal4/P{w[+mC]=UAS-TeTxLC.tnt}E2* | *Canton-S* | *Canton-S* | 18, 14 |
| | *w[1118];+/P{w[+mC]=UAS-TeTxLC.tnt}E2;+/Or65c-Gal4* | *Canton-S* | *Canton-S* | 20, 18 |
| | *w[1118];Or67d-Gal4/P{w[+mC]=UAS-TeTxLC.tnt}E2* | *Canton-S* | *Canton-S* | 15, 19 |
| | *w[1118];Or83c-Gal4/P{w[+mC]=UAS-TeTxLC.tnt}E2* | *Canton-S* | *Canton-S* | 20, 17 |
| | *w[1118];Or88a-Gal4/P{w[+mC]=UAS-TeTxLC.tnt}E2* | *Canton-S* | *Canton-S* | 21, 19 |

| | | | | |
|---|---|---|---|---|
| Fig. S2 | | | | |
| Fig. S2A | *w[1118]* | *Canton-S* | | 26, 14, 18, 13 |
| Fig. S2B | *w[1118]* | *Canton-S* | | 22, 14, 12, 12 |

| | | | | |
|---|---|---|---|---|
| Fig. S3 | | | | |
| Fig. S3A | *w[1118]* | *Canton-S* | | 33 |
| | *w[1118]* | *Canton-S* | *+;PromE(800)-Gal4/UAS-Tra* | 36 |
| | *w[1118]* | *Canton-S* | *+;PromE(800)-Gal4/UAS-Tra-RNAi* | 17 |
| Fig. S3B | *w[1118]* | *Canton-S* | | 20 |
| | *w[1118]* | *Canton-S* | *Drosophila melanogaster* | 23 |
| | *w[1118]* | *Canton-S* | *Drosophila simulans* | 21 |
| | *w[1118]* | *Canton-S* | *Drosophila sechellia* | 20 |
| | *w[1118]* | *Canton-S* | *Drosophila erecta* | 19 |
| | *w[1118]* | *Canton-S* | *Drosophila yakuba* | 21 |

| | | | | |
|---|---|---|---|---|
| Fig. S4 | *w[1118]* | *Canton-S* | | 18, 18, 14, 17, 15, 13 |

| | | | | |
|---|---|---|---|---|
| Fig. S5 | | | | |
| Fig. S5A | *w[1118]* | *Canton-S* | | 17, 17, 18, 18, 17, 18 |
| Fig. S5B | *w[1118]* | *Canton-S* | | 15, 15, 15, 16, 14, 15 |
| Fig. S5C | *w[1118]* | *Canton-S* | *Canton-S* | 21, 18 |
| | *w[1118]/ppk23-* | *Canton-S* | *Canton-S* | 17, 21 |
| | *ppk23-* | *Canton-S* | *Canton-S* | 13, 15 |
| Fig. S5D | *w[1118]* | *Canton-S* | *Canton-S* | 18, 14 |

| | | | | |
|---|---|---|---|---|
| | *w[1118]/ppk28-* | *Canton-S* | *Canton-S* | 23, 25 |
| | *ppk28-* | *Canton-S* | *Canton-S* | 22, 17 |
| Fig. S5E | *w[1118]* | *Canton-S* | *Canton-S* | 17, 14 |
| | *w[1118];ppk29-/+* | *Canton-S* | *Canton-S* | 19, 20 |
| | *ppk29-* | *Canton-S* | *Canton-S* | 16, 17 |

| | | | | |
|---|---|---|---|---|
| Fig. S6A-C | *w[1118];R52G04-AD/UAS-myrGFP;Dsx-DBD/UAS-myrGFP* | | | |
| Fig. S6D | *w[1118];R52G04-AD/+;Dsx-DBD/UAS-GtACR1* | *Canton-S* | | 82, 60 |

| | | | | |
|---|---|---|---|---|
| Fig. S7A | *w[1118];UAS-FLP/+; GMR71G01-Gal4, CRE-F-Luc/+* | | | 8, 8, 8, 12, 8, 4 |
| Fig. S7B | *w[1118];UAS-FLP/+; GMR71G01-Gal4, CRE-F-Luc/+* | | | 8, 8, 8, 12, 8, 4 |

| | | | | |
|---|---|---|---|---|
| Fig. S8 | *w[1118];UAS-FLP/+; GMR71G01-Gal4, CRE-F-Luc/+* | | | 12, 12, 12 |
| | *w[1118]; R52G04-AD/+;UAS-FLP, CRE-F-Luc/Dsx-DBD* | | | 12, 12, 12 |
| | *w[1118];;UAS-FLP, CRE-F-Luc/Dh44-pC1 (Dsx-DBD, Dh44A-AD)-GAL4* | | | 12, 12, 12 |
| | *w[1118]; VT25602-AD/+;UAS-FLP, CRE-F-Luc/VT2064-DBD* | | | 10, 10, 10 |

| | | | | |
|---|---|---|---|---|
| Fig. S9 | *w[1118]/UAS-Dcr2;;GMR71G01-Gal4/+* | *Canton-S* | *Canton-S* | 27, 26 |
| | *w[1118];UAS-Dh44R1-RNAi/+; UAS-Dh44R2-RNAi/+* | *Canton-S* | *Canton-S* | 18, 17 |
| | *w[1118]/UAS-Dicer2;UAS-Dh44R1-RNAi1/+; GMR71G01-GAL4/UAS-Dh44R2-RNAi2* | *Canton-S* | *Canton-S* | 35, 30 |

**Table. S1. The genotypes of the females and males, as well as the number of control and test females in each figure presented in this study**
